# CLM-X: A multimodal single-cell foundation model with flexible multi-way Transformer for unified scRNA-seq and scATAC-seq analysis

**DOI:** 10.64898/2026.02.17.704943

**Authors:** Bowen Li, Ziqiang Liu, Zhen Wang, Zhenyu Xu, Yantao Li, Chulin Sha, Xiaolin Li

## Abstract

Advances in single-cell multimodal profiling have enabled a more systematic analysis of cellular biology, yet the rapid accumulation of large-scale, heterogeneous datasets poses substantial challenges for integrative analysis. Recently, Transformer-based cell language models (CLMs) are becoming powerful foundational tools for learning transferable cell representations from unimodal single-cell datasets. However, a unified and flexible multimodal foundation model for joint modeling of scRNA-seq and scATAC-seq datasets remains underexplored. Here, we present CLM-X, a multimodal single-cell foundation model built on multiway Transformer architecture. CLM-X employs a harmonized tokenization design together with a stage-wise masked reconstruction pretraining strategy, enabling unified modeling of RNA-only, ATAC-only, and paired RNA–ATAC input within a single Transformer-based framework. We pretrain CLM-X on million-scale unimodal and multimodal datasets, and systematically evaluate its transferability on five downstream tasks including batch correction, modality integration, cross-modal translation, cell type annotation, and perturbation prediction. Across comprehensive benchmarks on 10 datasets, CLM-X consistently outperforms existing multimodal methods and unimodal foundation models, with particularly clear advantages in RNA–ATAC cross-modal translation and genetic-perturbation-response prediction. Overall, CLM-X establishes a unified and flexible multimodal foundation model for integrative analysis of scRNA-seq and scATAC-seq datasets, advancing a more robust, comprehensive, and biological interpretable single-cell analysis beyond current multimodal fusion approaches and unimodal foundation models.

## 1. Introduction

Cells are the fundamental structural and functional units of life. Recent advances in single-cell multimodal technologies have revolutionized our ability to profile individual cells across multiple molecular layers at unprecedented scale and resolution [1, 2]. By jointly measuring diverse molecular omics — transcriptomics, epigenomics, proteomics, and others — these technologies enable a more holistic view of cellular functions and regulatory states [3, 4]. Meanwhile, multimodal single-cell datasets are being generated at an ever-increasing speed, particularly accelerated by large atlas initiatives such as The Human Cell Atlas (HCA) [5] and the Human BioMolecular Atlas Program (HuBMAP) [6]. These integrative datasets provide great opportunities to investigate complex problems in molecular cell biology, such as how cells maintain homeostasis, transit between functional states, and respond to perturbations [7, 8, 9]. However, such large-scale, high-dimensional, and structurally heterogeneous datasets pose substantial analytical challenges. On one hand, the intrinsic diversity of molecular modalities and the complex, often nonlinear relationships among them make multimodal data integration inherently difficult. Different omics exhibit distinct technical biases and noise effects, further complicating the analysis [10]. On the other hand, the explosive accumulation of single-cell datasets has exposed the limitations of existing methods, most of which are designed for specific tasks and lack the scalability required to effectively handle datasets at the million-cell scale. Together, these challenges underscore the urgent need for new computational paradigms that can effectively leverage large-scale, multimodal single-cell datasets, capturing cross-modal interrelations while remaining scalable and flexible, thereby enabling deeper and more systematic biological discovery [11, 12].

A vast number of machine learning methods have been developed to integrate single-cell multimodal data, such as MultiVI [13], Multigrate [14], MIRA [15], scJoint [16], scBridge [17], scMoMaT [18], and etc. These methods typically employ joint embedding strategies to align multimodal profiles in a shared latent space, and have substantially advanced the integration analysis of multimodal single-cell datasets. Their performance has been systematically evaluated in several benchmarking studies [19, 20] across diverse analytical tasks and settings. Despite these advances, these models face several fundamental limitations. First, most existing methods involve modality-specific preprocessing and normalization steps to reduce input dimensionality for computational efficiency, which can introduce subjective biases and complicate integrative analysis. Moreover, many of these methods are designed for specific tasks and are trained on relatively limited datasets, preventing them from fully exploiting the millions of cells available in atlas-scale single-cell resources. As a result, they often struggle to learn consistent, transferable representations that reflect conservative biological patterns.

Recently, the Cell Language Models (CLMs) or Foundation Models paradigm — inspired by large language models in natural language processing — has emerged as a powerful framework and demonstrated substantial potential in capturing cellular biological patterns. These foundation models are typically pretrained on large-scale single-cell datasets and subsequently adapted to diverse downstream tasks via transfer learning. For scRNA-seq data analysis, models such as Geneformer [21], scGPT [22], and scFoundation [23] have shown remarkable cell representational ability and transferability, enabling robust performance across a wide range of downstream tasks with fine-tuning. More recently, this paradigm has been extended to scATAC-seq data analysis, models such as GET [24], EpiAgent [25], and CLM-Access [26] are pretrained on large scATAC-seq datasets to learn the regulatory logic of chromatin states and support further related analysis. Together, these successes highlight the potential of unimodal foundation models to extract conserved biological knowledge from large-scale single-cell datasets.

This naturally leads to the expectation that a foundation model trained on large single-cell multimodal datasets could further enhance representation learning by leveraging complementary transcriptional and epigenetic information [27, 28]. However, only a limited number of studies—most notably scCLIP [29] and SCARF [30]—have attempted to jointly model scRNA-seq and scATAC-seq profiles in a cell foundational model manner. Both models rely primarily on contrastive pretraining objectives, which align matched RNA–ATAC pairs by pulling paired instances together and pushing unpaired instances apart. While effective for cross-modal alignment, contrastive objectives are often limited in their ability to capture complementary information across modalities. More importantly, contrastive learning frameworks are typically tailored to specific pairs of modalities. Consequently, their performances are constrained by the available paired data, and lack the flexibility in handling unpaired or more than two modalities. Therefore, the field still lacks of a flexible multimodal foundation model that can unify scRNA-seq and scATAC-seq analysis, demonstrating consistent performance gains across diverse downstream tasks, while remaining agnostic to input technologies and free from manual annotation requirements.

To meet this unfulfilled need, we propose CLM-X, a multimodal foundation model for unified integration of scRNA-seq and scATAC-seq data(Fig. 1a-c). CLM-X is built upon the BEiT-3 architecture [31]. BEiT-3 adopts multiway Transformer as a unified backbone and masked data modeling as a unified pretraining objective, supporting a general-purpose multimodal foundation model framework. Building on this architecture, CLM-X consists of three Transformer branches: two unimodal branches for scRNA-seq and scATAC-seq data, respectively, and a multimodal branch for paired scRNA–scATAC data which learns RNA-ATAC interaction. This allows CLM-X to exploit the abundant single-modality datasets while reducing the dependence on large paired datasets. Specifically, the multimodal branch are pretrained on paired scRNA–scATAC datasets via a cross-modal conditional mask and reconstruction objective, encouraging the model to capture alignments and complementarities between two modalities. We also designe a harmonized tokenization for scRNA-seq and scATAC-seq data in CLM-X. Both modalities are represented as uniform token sequences with a fixed-length of 2,000. This tokenization scheme enables seamless integration within a shared multi-way Transformer backbone. Altogether, CLM-X provides a flexible and generalizable foundation model for scRNA-seq and scATAC-seq integration that can naturally scale to heterogeneous unpaired single-cell multimodal datasets.

**Fig. 1:**
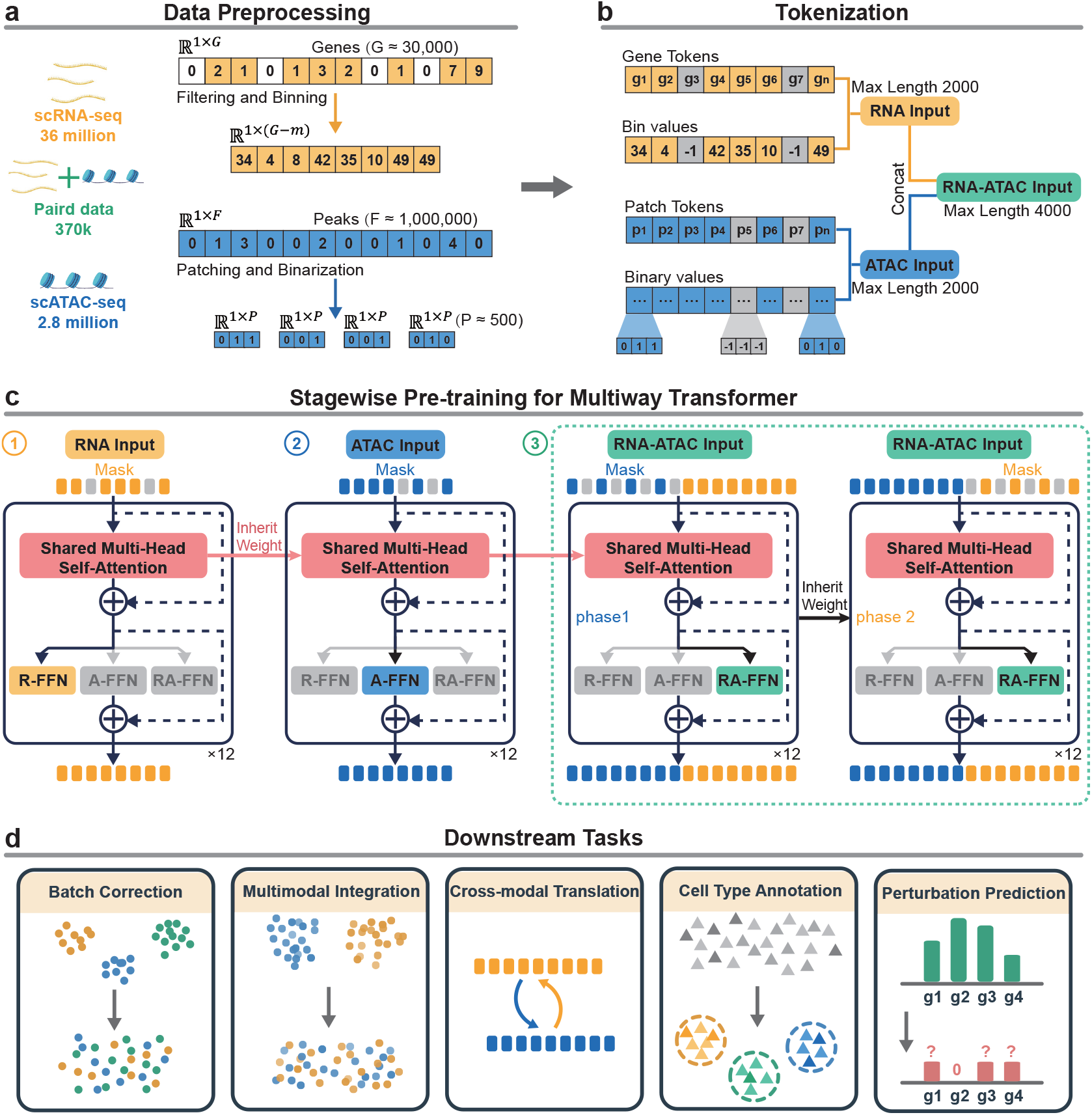
CLM-X workflow. **a**, Pretraining data and modality-specific preprocessing. CLM-X is pretrained on scRNA-seq (36 million cells), scATAC-seq (2.8 million cells), and paired RNA–ATAC data (370k cells). scRNA-seq profiles span ∼30,000 genes; non-zero counts are discretized into rank-preserving bins. scATAC-seq peak counts (∼ 1,000,000 peaks) are binarized, ordered by genomic coordinates, and grouped into fixed-length, genome-ordered patches. **b**, Tokenization and input packing. Cells are encoded as gene tokens with binned expression values (RNA) or genome-ordered patch tokens with within-patch binary accessibility vectors (ATAC). At each position, the input is the element-wise sum of token and value embeddings. Unimodal sequences are padded/truncated to 2,000 tokens; paired RNA–ATAC inputs concatenate RNA and ATAC into one context window (up to 4,000 tokens) for cross-modal interaction via shared self-attention. **c**, Multiway Transformer backbone and stage-wise pretraining. CLM-X uses a multiway Transformer with shared MHSA and input-type-specific FFN experts (R-FFN, A-FFN, RA-FFN). The model is pretrained with a unified masked reconstruction objective in a stage-wise strategy with parameter inheritance: (1) RNA-only masked RNA values reconstruction; (2) ATAC-only masked peak values reconstruction while inheriting shared MHSA weights from stage (1); and (3) paired RNA–ATAC fusion with two-phase conditional reconstruction (mask ATAC and reconstruct ATAC given visible RNA, then mask RNA and reconstruct RNA given visible ATAC), enabling bidirectional cross-modal translation and stable fusion training. **d**, Downstream applications include batch correction, multimodal interation, cross-modal translation, cell type annotation, and perturbation prediction.

We pretrain CLM-X on three large datasets comprising 36 million scRNA-seq cells, 2.8 million scATAC-seq cells, and 370 thousand paired scRNA-scATAC cells. Based on this pretrained model, we provide task-specific fine-tuning pipelines for five widely used single-cell analysis tasks (Fig. 1d)including: batch correction, multimodal integration, RNA–ATAC cross-modal translation, cell type annotation, and gene perturbation prediction. We evaluate the transferability of CLM-X on a comprehensive benchmark of 10 datasets, comparing it against diverse state-of-the-art multimodal integration methods as well as scRNA-seq–based foundation models. CLM-X shows consistently improved performance across all tasks. In particular, CLM-X achieves notable gains in RNA–ATAC cross-modal translation, where it supports unfiltered-input modeling and full-scale cross-modal prediction, surpassing the capabilities of current methods. CLM-X also outperforms task-specific models and unimodal foundation models in cell type annotation and gene perturbation prediction tasks. These results indicate that CLM-X provide a unified and scalable multimodal foundation model for integrative analysis of scRNA-seq and scATAC-seq data, supporting more robust and comprehensive single-cell analysis beyond unimodal and task-specific frameworks.

## 2 Results

### 2.1 Overview of CLM-X

Single-cell multimodal foundation models hold great promise for the integration of large-scale multi-omics datasets. However, compared to unimodal foundation models, the development of multimodal foundation models faces substantially higher challenges in flexibility, scalability, and generalizability. First, since different omics modalities exhibit profound heterogeneity in data structure and noise characteristics, designing a unified input form that can retain full modality-specific information without heavy manual filtering or data transformation is a key challenge. Second, a general-purpose pretraining architecture is required to enable modality-agnostic deep fusion and information augmentation, meanwhile remaining scalable and easily transferable to new tasks or modalities. Third, as high-quality paired multimodal data is still scarce, how to maximize the utility of existing unimodal data and minimize the dependence on paired multimodal data is also a practical concern.

To systematically address these challenges, we develop CLM-X—a multimodal foundation model for the unified analysis of scRNA-seq and scATAC-seq data. The overview design involved with tokenization, pretraining, and fine-tuning is illustrated in Fig. 1. We collect over 36 million scRNA-seq data that covers cells of diverse human tissue from CellXGene[32], 2.8 million scATAC-seq data assembled in our previous work CLM-Access [26], and 370 thousand simulated scRNA-seq and scATAC-seq paired data from scCLIP [29]. We introduce a unified representation scheme consists of a token embedding and a value embedding (Fig. 1a), which enables both modalities to be modeled within a consistent input space. Specifically, for scRNA-seq data, CLM-X adopts a scGPT-inspired tokenization strategy, in which each cell is represented as a sequence of gene tokens paired with binned expression values. For scATAC-seq data, the ultra-high dimensionality of 10^6^ peaks are contiguously grouped into patches and binarized within each patch, and cells are represented as patch tokens paired with binary vectors the same way used in CLM-Access [26]. Each modality is padded or truncated to a maximum of 2,000 tokens. For paired RNA–ATAC cells, RNA and ATAC token sequences are concatenated along the sequence dimension to form a single context window of up to 4,000 tokens (Fig. 1b), allowing cross-modality information to interact directly through self-attention within a shared context.

To enable generalizable multimodal pretraining, CLM-X adopts BEiT-3 architecture[31], which applies a “shared attention with modality-specific feed-forward networks” backbone to achieve deep multimodal fusion while preserving modality-specific representations. This allows CLM-X to accommodate RNA-only, ATAC-only, and paired RNA–ATAC inputs within a single framework. In addition, CLM-X adapts masked data modeling for multimodal pretraining, coupled with a stage-wise pretraining strategy. (Fig. 1c). Pretraining proceeds by first performing unimodal masked reconstruction on scRNA-seq and scATAC-seq datasets sequentially to initialize shared attention parameters, followed by cross-modal conditional reconstruction on paired data to learn multimodal alignment and complementary information. During paired-data pretraining, RNA and ATAC token sequences are fused into a shared context and optimized by a two-phase masked reconstruction scheme. In phase I, masks are applied exclusively to ATAC tokens and the model reconstructs chromatin accessibility conditioned on the RNA context. In phase II, masking is applied to RNA tokens while reconstruction is conditioned on the ATAC context, with parameters inherited from the first phase (See details in Methods). This bidirectional conditional cross-modal reconstruction enables the model to capture multimodal dependencies in both directions, while improving training stability and fusion comprehensiveness under limited paired data availability.

After pretraining, we fine-tune CLM-X on five downstream tasks(Fig. 1d): batch-effect correction, RNA–ATAC modality integration, bidirectional cross-modal translation, multimodal cell type annotation, and perturbation-response prediction, by introducing lightweight task-specific heads, while keeping the core encoder architecture unchanged. Downstream objectives are implemented through minimal task-specific decoders or classifiers operating on shared cell embeddings, thereby leveraging the multimodal alignment and cross-modal dependencies established during pretraining (implementation details in Methods 4.3). Collectively, CLM-X establishes a unified and scalable framework for multimodal single-cell analysis, demonstrating how a single pretrained multimodal encoder can be systematically adapted to heterogeneous downstream tasks.

### 2.2 CLM-X Removes Batch Effects While Preserving Biological Signals

Batch effects among different platforms, experiments and studies can obscure the true structure of cellular states [33, 34]. To systematically assess batch correction and dataset integration, we evaluate on three representative multi-batch benchmarks: PBMC (Datasets 1–4; including granulocyte-depleted and unsorted samples; 4 batches), PBMC (Datasets 6–7; 2 batches), and BMMC (Dataset 5; multi-donor, cross-center; 13 batches). We use NMI to quantify preservation of biological structure and bASW to quantify batch mixing, and report their arithmetic mean as the overall score. We evaluate the zero-shot and fine-tuned setting of CLM-X. CLM-X is compared against four multimodal integration including MultiVI [13], scMoMaT [18], Multigrate [14], and MIRA [15] as well as the RNA-unimodal foundation model scGPT.

Across the three benchmarks, CLM-X consistently achieves the strongest balance between biological structure preservation and batch mixing. It attains the highest mean NMI, bASW and overall score on each dataset, demonstrating stable integration performance across varying degrees of technical complexity(Fig. 2a–c; Supplementary Table S1). On PBMC (Datasets 1–4), CLM-X preserved clear cell type structure while effectively mixing batches (Fig. 2d). Similar trends are observed on PBMC (Datasets 6–7) S1) and the more heterogeneous BMMC cohort(Fig. 2e), where CLM-X maintains robust performance despite multi-donor and cross-center variation. Overall, CLM-X improves the overall batch-correction score by 5.9%–35.0% compared with multimodal baselines (Fig. 2c). Importantly, zero-shot CLM-X produces meaningful embeddings without any task-specific training, indicating that multimodal pretraining alone captures substantial batch-invariant biological structure. Compared with the RNA-only foundation model scGPT, CLM-X achieves higher overall scores across all datasets, with an average relative improvement of 11.6%. These results suggest that multimodal pretraining followed by paired multiome adaptation yields cell-state representations that more effectively reconcile biological conservation with batch removal under substantial technical heterogeneity.

**Fig. 2:**
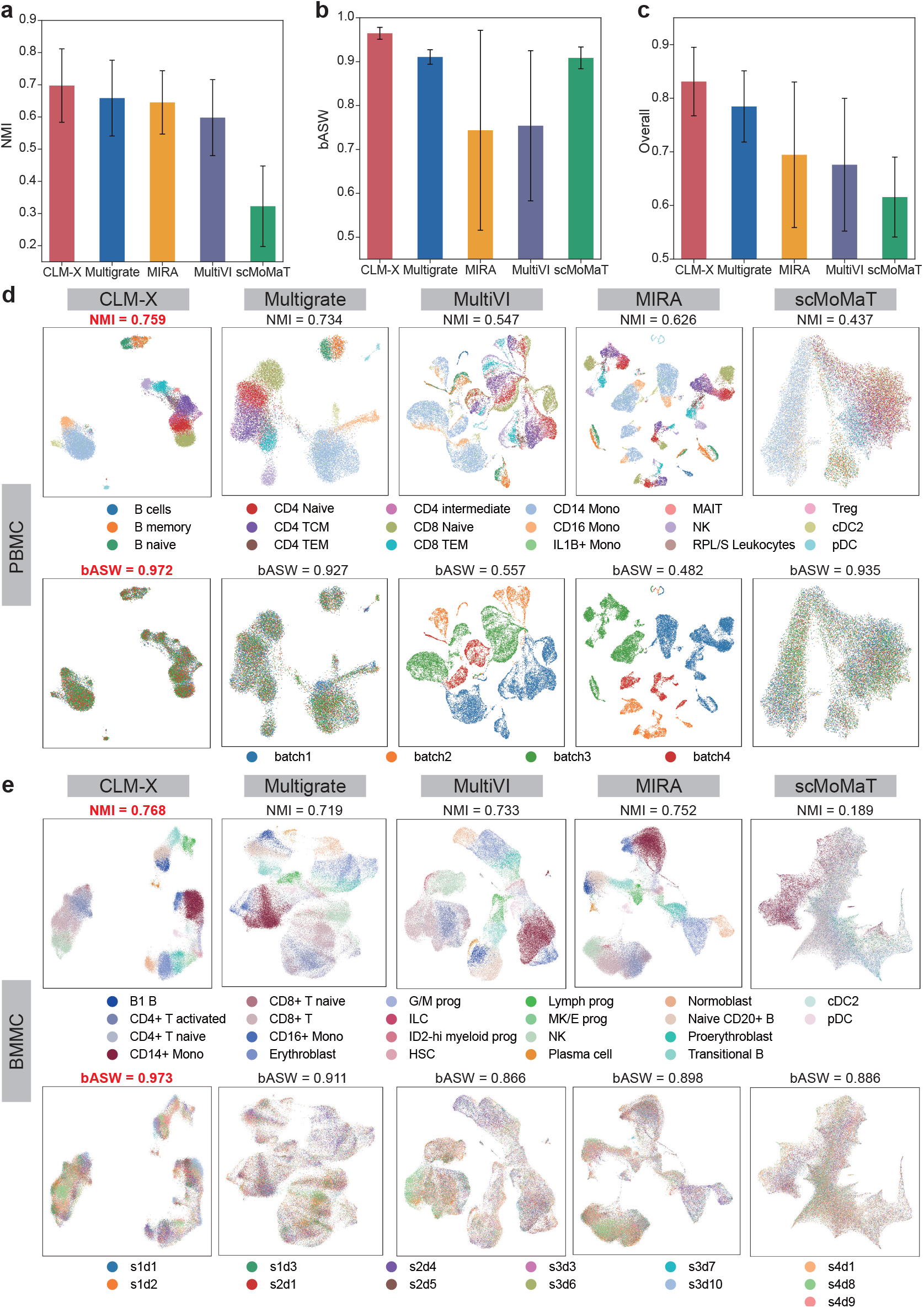
Results of batch-effect correction. **a–c**, Bar plots summarizing batch-correction performance across the three benchmarks (PBMC datasets 1–4; BMMC dataset 5; and PBMC datasets 6–7). Bars and error bars denote mean ±s.d. across benchmarks. **a**, NMI measuring preservation of cell type structure. **b**, bASW measuring batch-effect removal. **c**, Overall score computed as the arithmetic mean of NMI and bASW. **d**, PBMC benchmark (datasets 1–4, 4 batches): UMAP visualization of integrated embeddings from CLM-X and multimodal baselines (Multigrate, MultiVI, MIRA and scMoMaT). For each method, embeddings are colored by cell type (top) and by batch (bottom). NMI and bASW are reported above the corresponding panels. **e**, BMMC benchmark (dataset 5, 13 batches) shown in the same format.

Collectively, CLM-X exhibits the most favorable NMI–bASW trade-off among all compared methods, achieving strong batch mixing while retaining well-resolved cellular identities. Such representation quality is essential for constructing integrated multimodal references across heterogeneous protocols and centers, reducing the risk that technical variation is misinterpreted as biological signal in downstream analyses.

### 2.3 CLM-X Enhances Cell Profiling by Integrating Expression and Chromatin Accessibility

Multimodal integration aims to fuse scRNA-seq and scATAC-seq from the same cells into a unified representation, leveraging the interpretability of RNA and the regulatory information captured by chromatin accessibility; however, the two modalities reside in heterogeneous feature spaces and exhibit markedly different sparsity/noise characteristics, making naive fusion prone to modality dominance or distortion of local neighborhood structure [3, 10, 11, 13]. We evaluate the integration performance on Datasets 1–7 (six PBMC multiome datasets and one BMMC dataset, spanning multiple platforms and batch configurations) using ARI/NMI (agreement between clustering and cell-type labels), cASW (cell-type separability) and cLISI (local consistency of cell-type neighborhoods), and aggregate them into an Overall fusion score. CLM-X addresses this task by fine-tuning to produce cell-level fused embeddings; following the benchmark-recommended protocol [19], we compare with MultiVI [13], SCOIT [35], Multigrate [14], and MIRA [15], and report CLM-X RNA-only and ATAC-only embeddings as ablations to quantify the gain from fusion.

As summarized in Fig. 3a, CLM-X achieves the highest aggregated fusion score (Overall = 0.730), outperforming all compared multimodal baselines (Supplementary Table S2). CLM-X leads in clustering agreement overall (ARI = 0.658, NMI = 0.716) while maintaining strong cell type neighborhood structure (cLISI = 0.977) and competitive separation (cASW = 0.569) (Supplementary Table S2). In the ARI–NMI summary (Fig. 3b), CLM-X occupies the top-right region (mean ± s.d. across datasets), indicating improved recovery of cell type structure. In the cASW–cLISI summary (Fig. 3c), CLM-X achieves consistently high local neighborhood consistency with comparable separation to the strongest baselines, suggesting that gains in clustering agreement are not obtained at the expense of local structure. Consistent with these trends, CLM-X attains the best overall performance with a 1.8%–16.4% relative improvement over multimodal baselines (Fig. 3d).

**Fig. 3:**
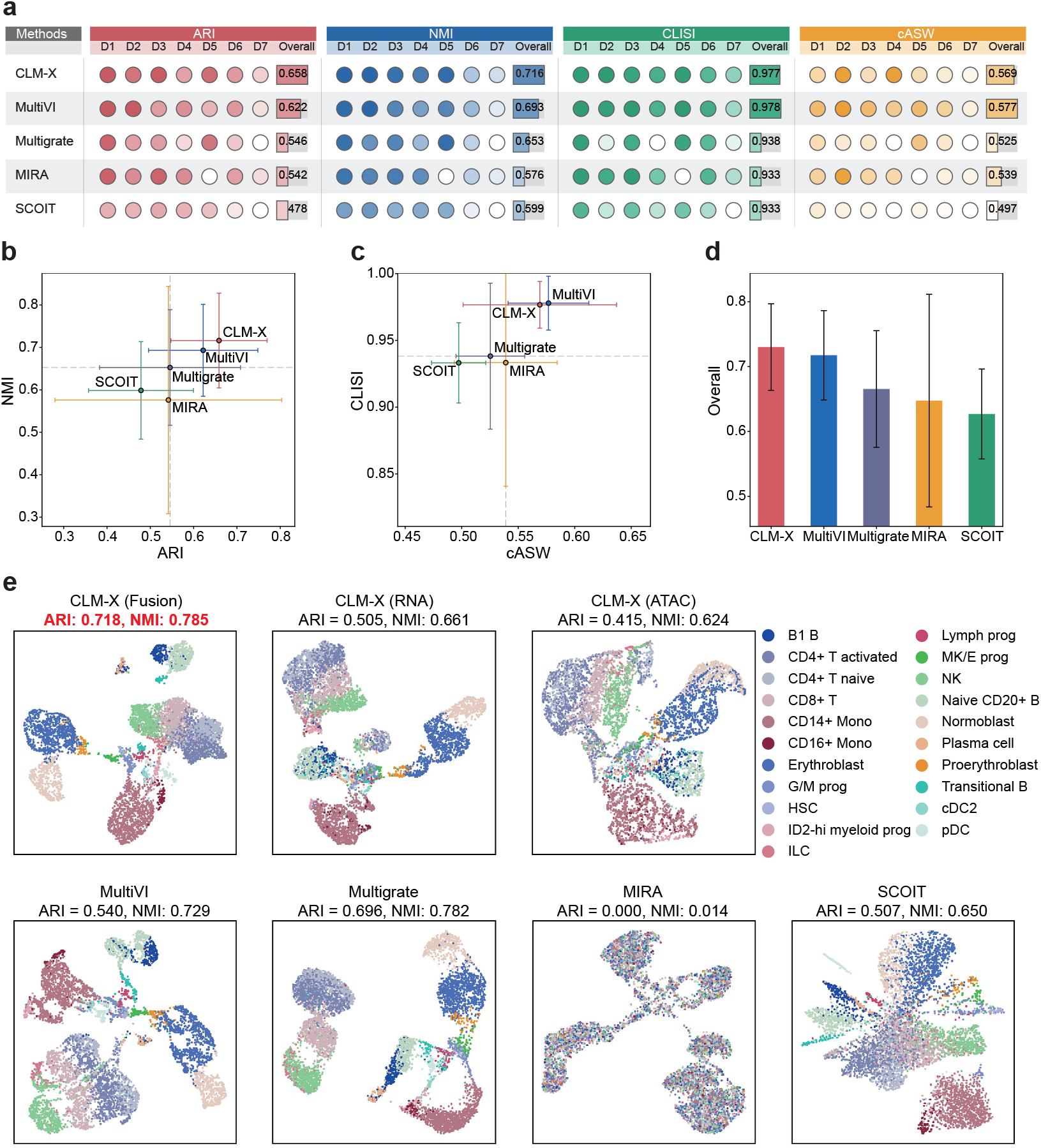
Modality fusion of RNA expression and chromatin accessibility. **a**, Bubble plot summarizing per-dataset (D1–D7) and overall performance for CLM-X and baseline methods (MultiVI, Multigrate, MIRA, and SCOIT) evaluated by ARI, NMI, cLISI and cASW (higher is better). **b**, Summary of clustering agreement with cell type annotations in the fused embedding, shown as ARI and NMI (mean ± s.d. across datasets). **c**, Summary of cell type structure in the fused embedding, shown as cASW and cLISI (mean ± s.d. across datasets). **d**, Overall fusion score aggregated from ARI, NMI, cASW, and cLISI across datasets (mean ± s.d.). **e**, UMAP visualization of dataset 5 embeddings colored by cell type, comparing CLM-X (fusion) with CLM-X unimodal embeddings (RNA-only and ATAC-only) and multimodal baselines; ARI and NMI are reported above each panel.

Qualitative visualization further corroborates the integration advantage: in Dataset 5 (Fig. 3e), CLM-X (Fusion) produces the clearest cell type organization (ARI = 0.718, NMI = 0.785), outperforming its RNA-only and ATAC-only embeddings. Among multimodal baselines, CLM-X also exceeds MultiVI and SCOIT, and slightly improves over Multigrate; in contrast, MIRA shows poor separation on this dataset. Detailed per-dataset results are provided in Supplementary Table S2 and Supplementary Figs. S2.

In general, CLM-X achieves the highest aggregated fusion score in Datasets 1–7, improving clustering agreement with cell type while maintaining strong local neighborhood consistency in fused embedding. The consistent advantage over RNA-only and ATAC-only ablations supports that CLM-X leverages complementary transcriptional and regulatory information in a balanced way, establishing that the fused embedding yields more informative cell representation than either modality alone.

### 2.4 CLM-X Excels at Quantitative Reconstruction of Accessibility and Expression

Cross-modality translation evaluates whether a model can infer an unobserved modality from an observed one, enabling missing-modality imputation and joint analyses [36, 13]; this task is challenging due to the mismatch between gene and peak feature spaces, the extreme sparsity of scATAC-seq, and the nonlinear, context-dependent coupling between accessibility and transcription [10]. For each dataset in Datasets 1–7, we randomly split cells into a training set comprising 80% of cells and a held-out test set comprising 20% of cells, using stratified sampling by cell-type labels to ensure matched label distributions between the two splits; we evaluate both RNA → ATAC and ATAC → RNA translation (with both all-gene and 2,000-HVG targets), reporting AUROC for accessibility prediction and PCC/RMSE for quantitative reconstruction fidelity (Adj. RMSE is used only for radar-plot visualization). CLM-X is fine-tuned for cross-modality translation to predict the target modality from the observed modality, thereby learning a transferable mapping between RNA expression and chromatin accessibility; we compare against BABEL [36], MultiVI [13] and CMAE [37] under the benchmark-recommended settings [19].

For the RNA → ATAC translation task over the full peak set, CLM-X yields the most accurate quantitative reconstruction, as evidenced by higher PCC and lower RMSE relative to all baseline approaches. In particular, CLM-X achieves the lowest RMSE on each of the seven benchmark datasets and the highest cross-dataset mean PCC (Fig. 4a; Supplementary Tables S3). For binary discrimination performance, CLM-X attains an AUROC comparable to MultiVI (0.959 vs. 0.968). Given this similar AUROC, the gain in PCC/RMSE is especially notable: CLM-X more faithfully recapitulates both peak-level accessibility magnitudes and cell-to-cell variability, rather than only the binary on/off pattern.

**Fig. 4:**
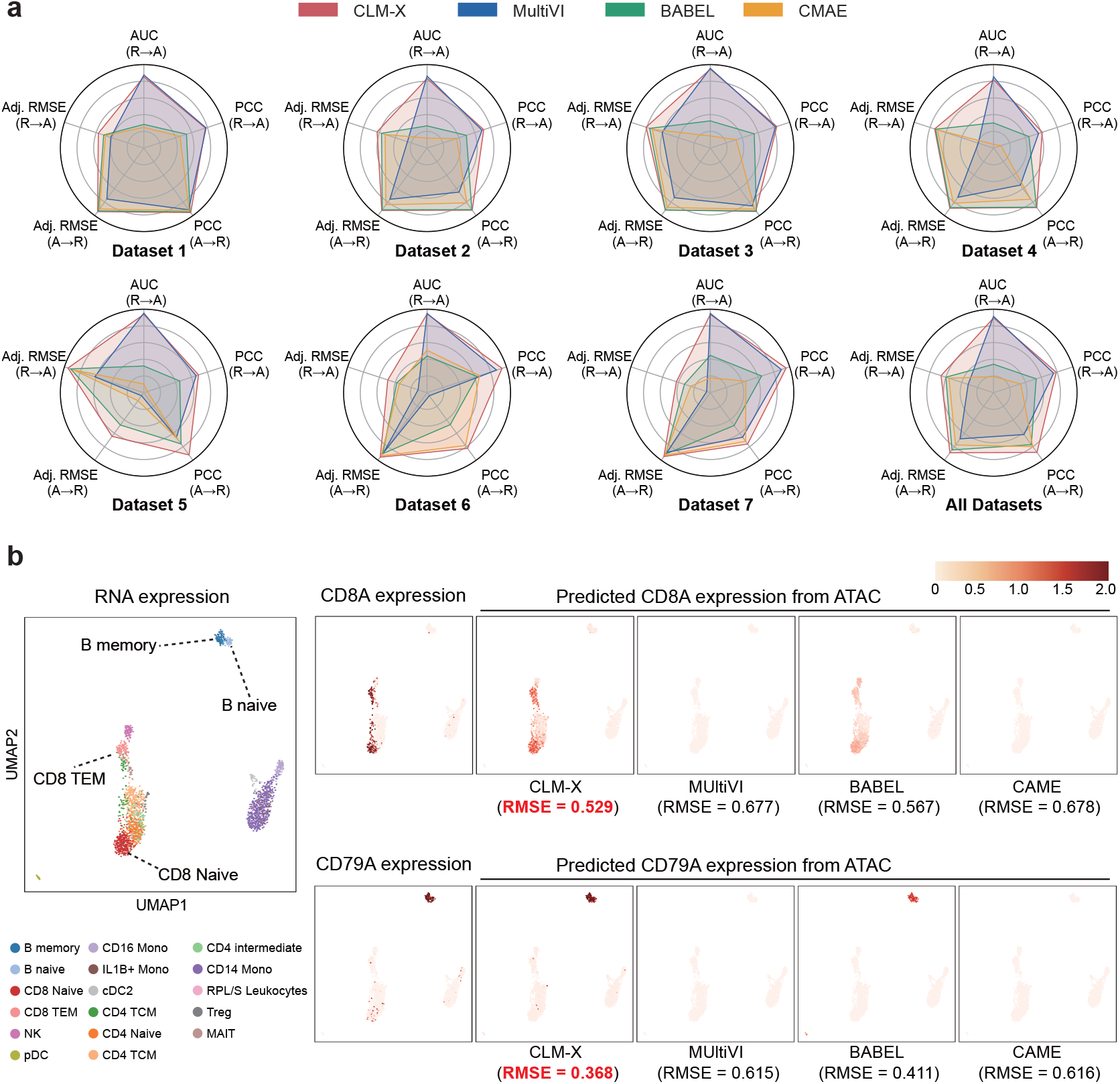
Cross-modal translation between RNA and ATAC. **a**, Radar plots summarizing cross-modal translation performance on seven datasets (datasets 1–7) and the aggregated result across datasets (All Datasets), comparing CLM-X with MultiVI, BABEL, and CMAE. Metrics include AUC for RNA → ATAC prediction, and PCC and adjusted RMSE (Adj. RMSE; min–max normalized and inverted for visualization) for both RNA → ATAC and ATAC → RNA prediction. **b**, Example from dataset 1: UMAP constructed from ground-truth RNA expression and annotated by cell type (left). Ground-truth expression of marker genes CD8A and CD79A (middle), and ATAC → RNA-predicted expression from each method (right), with per-gene RMSE reported below each prediction panel.

For the ATAC → RNA task using all genes, CLM-X again delivers the strongest overall performance: it achieves the highest PCC on six of the seven datasets and the lowest RMSE on all seven datasets (Fig. 4a; Supplementary Tables S3). Among baselines, CMAE ranks second in mean PCC (0.787 ± 0.074) but exhibits substantial cross-dataset variability in RMSE (3.333 ± 3.394), comparable to MultiVI (4.395 ± 3.346); differences in expression scale across datasets can amplify RMSE instability in cross-dataset comparisons. In contrast, CLM-X shows consistently low errors across datasets, indicating stronger robustness and generalization. When restricting ATAC → RNA to 2,000 HVGs, CLM-X achieves the highest PCC while simultaneously attaining the lowest RMSE on all seven datasets (Supplementary Tables S3), demonstrating improved quantitative fidelity on the core genes driving biological variation.

To qualitatively assess whether CLM-X preserves cell type-specific expression patterns in held-out cells, we computed a UMAP embedding of the RNA test set in Dataset 1 using ground-truth expression (Fig. 4b) and overlaid the predicted ATAC → RNA expression values. Using CD8A and CD79A as marker genes for CD8^+^ T cells and B cells, respectively, CLM-X predictions recapitulate spatial distributions and expression patterns that closely match the ground truth. Together, these results indicate that CLM-X not only improves global translation metrics, but also preserves biologically meaningful, cell type-specific signals.

Overall, across seven datasets and both translation directions, CLM-X delivers consistently improved quantitative reconstruction performance, and preserves cell type-specific marker patterns in held-out cells. These findings suggest that CLM-X learns transferable transcription–chromatin couplings that enable precise cross-modal prediction and provide a foundation for further dissecting transcriptional programs and their epigenetic regulatory mechanisms.

### 2.5 CLM-X Delivers Accurate and Stable Multimodal Cell Type Annotation

Cell type annotation is fundamental for characterizing single-cell composition and heterogeneity [38]; in paired RNA–ATAC settings, a practical goal is to annotate cells based on fused representations that exploit complementary information from both modalities while remaining robust to ATAC sparsity and cross-dataset variation. We evaluate on four matched 10x Genomics PBMC multiome datasets (Datasets 1–4; same donor; granulocyte-depleted vs. unsorted; two capture scales), and within each dataset we randomly split cells at the cell level into a training set comprising 80% of cells and a held-out test set comprising 20% of cells, using stratified sampling by cell-type labels to ensure matched label distributions between the two splits; we report Accuracy and Macro F1. CLM-X performs supervised fine-tuning for classification using fused cell embeddings, and we compare with Seurat (WNN) as well as multimodal variants of scGPT, scBridge, and scJoint (implementation and adaptation details are provided in Methods).

As shown in Fig. 5a, CLM-X (Fusion) provides the best overall performance across datasets, achieving the highest mean Macro F1 (88.24 ± 2.35%) together with consistently strong Accuracy (91.88 ± 1.19%) (Table S4). Fusion yields clear gains over unimodal variants: Macro F1 increases from 86.86 ± 1.77% (RNA-only) and 67.45 ± 9.01% (ATAC-only) to 88.24 ± 2.35%, indicating that CLM-X effectively benefits from complementary information when both modalities are available (Table S4). Relative to external baselines, CLM-X (Fusion) improves average Macro F1 compared with Seurat (WNN) (85.44 ± 6.67%) and substantially outperforms scGPT (74.94 ± 7.75%) and scJoint (58.14 ± 9.48%), while also exhibiting smaller cross-dataset variability than most baselines (Table S4), indicating stronger stability across matched datasets.

**Fig. 5:**
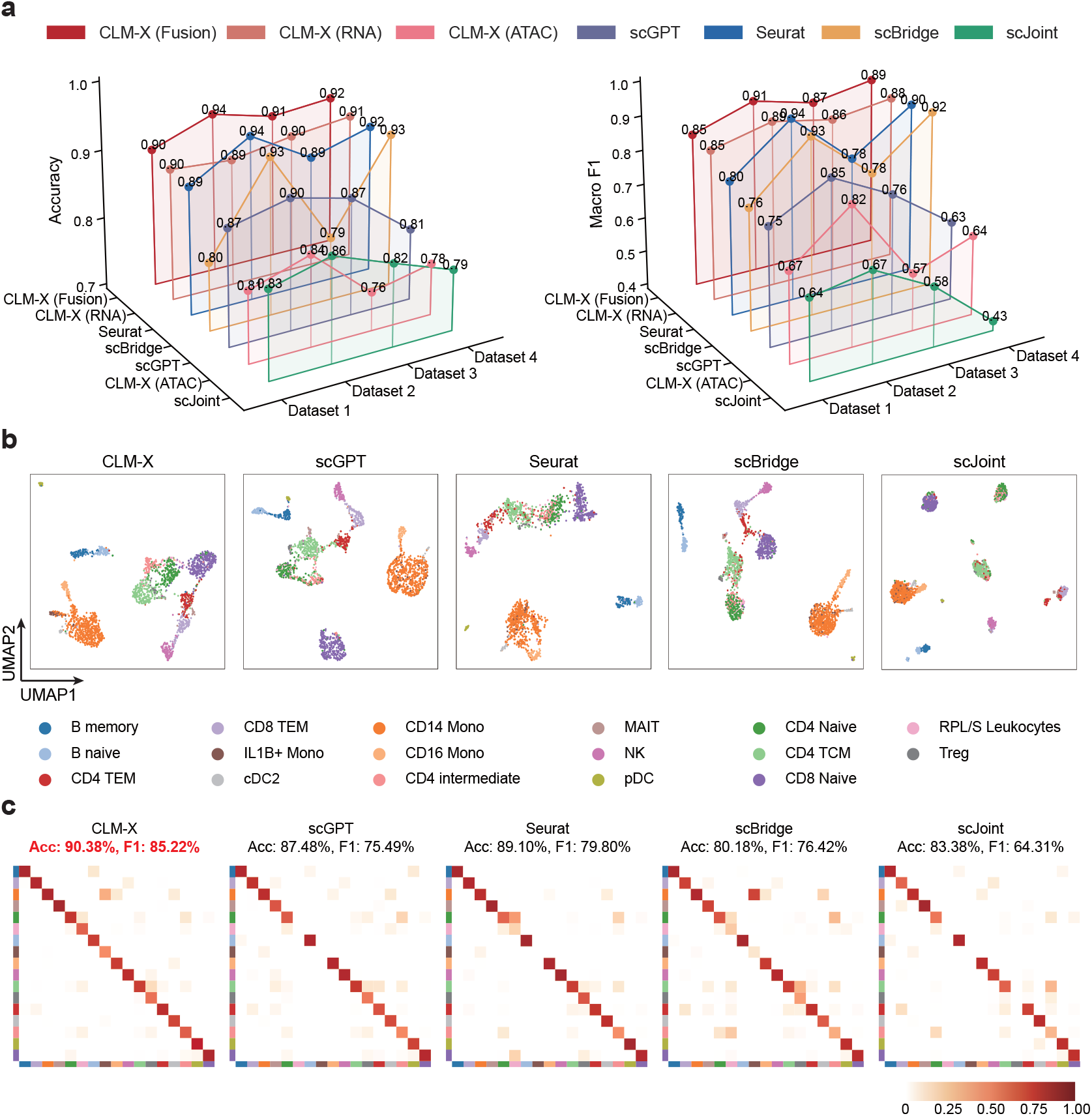
Multimodal cell type annotation results. **a**, Per-dataset annotation performance across four PBMC multiome datasets, reported as Accuracy (left) and Macro F1 (right) for CLM-X (Fusion), CLM-X (RNA), CLM-X (ATAC), scGPT, Seurat, scBridge, and scJoint. **b**, UMAP visualization of Dataset 1 ATAC-derived embeddings colored by predicted cell type labels for CLM-X and representative baselines. **c**, Confusion matrices on Dataset 1 comparing predicted versus reference cell type labels; Accuracy (Acc.) and Macro F1 are reported above each matrix.

On Dataset 1, UMAP visualizations of fused embeddings colored by predicted labels (Fig. 5b) show that CLM-X produces a more coherent and well-separated cell type structure than representative baselines. Confusion matrices (Fig. 5c) further show reduced off-diagonal errors, especially for challenging populations. Two notable rare/difficult classes are IL1B+ Mono and RPL/S Leukocytes, which often sit close to abundant neighboring types and can be driven by transient activation/stress programs [38]. In class-wise analysis (Table S5), CLM-X achieves substantially higher recall on IL1B+ Mono (82.35%) than scBridge (70.59%), whereas scGPT, scJoint, and Seurat fail to recover this class (0% recall) and instead collapse it into CD14 Mono. CLM-X also improves precision for IL1B+ Mono (57.14%) compared with scBridge (15.09%), indicating fewer CD14 Mono are incorrectly assigned to the IL1B+ subtype. For RPL/S Leukocytes, CLM-X reaches 39.29% recall, far exceeding Seurat (3.57%) and matching scBridge (39.29%), while achieving much higher precision (73.33% vs. 28.21%). Additional qualitative results for Dataset 2–4 are provided in Supplementary Figs. S1.

In multimodal cell type annotation on the evaluated multiome datasets, CLM-X delivers the highest overall accuracy and the most consistent performance, with particularly clear improvements on closely related and confusable populations. This suggests that the fused embedding provides more reliable decisions. These results validate the effectiveness of our fusion framework and suggest that it can be readily extended to other multimodal task scenarios.

### 2.6 CLM-X Predicts Transcriptomic Responses to Unseen Genetic Perturbations

Gene-perturbation prediction aims to forecast how perturbations alter cellular transcriptional states, supporting genefunction discovery, regulatory-dependency identification, and in silico screening; a central challenge is extrapolating to target-gene perturbations unseen during training. We evaluate on three Perturb-seq datasets using a held-out perturbation split to ensure that test perturbation conditions never appear in training: Adamson (K562; 86 single-gene perturbations), replogle k562 essential (K562; 412 perturbations after filtering), and replogle rpe1 essential (RPE1; 652 perturbations after filtering). Using the perturbation-induced differential expression vector (relative to control) as the signature, we report DE20 Pearson (correlation on the top-20 true DE genes) and ! Pearson (genome-wide correlation), and additionally provide top-*k* DE gene recovery statistics to assess signature fidelity. This task is conducted in an RNA-only fine-tuning setting, where CLM-X learns transferable perturbation responses by regressing the post-perturbation expression profile conditioned on the perturbation; we compare against GEARS [39] and scGPT [22] under their recommended settings.

Across all three datasets, CLM-X achieves the highest DE20 Pearson, improving over the best baseline by +0.0029 (Adamson), +0.0196 (Replogle K562 Essential), and +0.1293 (Replogle RPE1 Essential) (Table S6). For ! Pearson, CLM-X outperforms baselines on the two Replogle datasets (+0.0332 on K562 and +0.0437 on RPE1 over the best baseline) and remains near-tied with the strongest baseline on Adamson (0.7423 vs. 0.7467 for GEARS), while substantially improving over scGPT. Figure 6a further shows that CLM-X yields higher overlap with ground-truth top-*k* DE genes and ranks true DE genes closer to the top of its predicted list across multiple *k* values, indicating more faithful recovery of both the responsive gene set and its ordering. Case studies on Adamson (Fig. 6b) illustrate that CLM-X captures the direction and relative magnitude of expression changes for top DE genes, with predicted distributions aligning with ground-truth mean shifts.

**Fig. 6:**
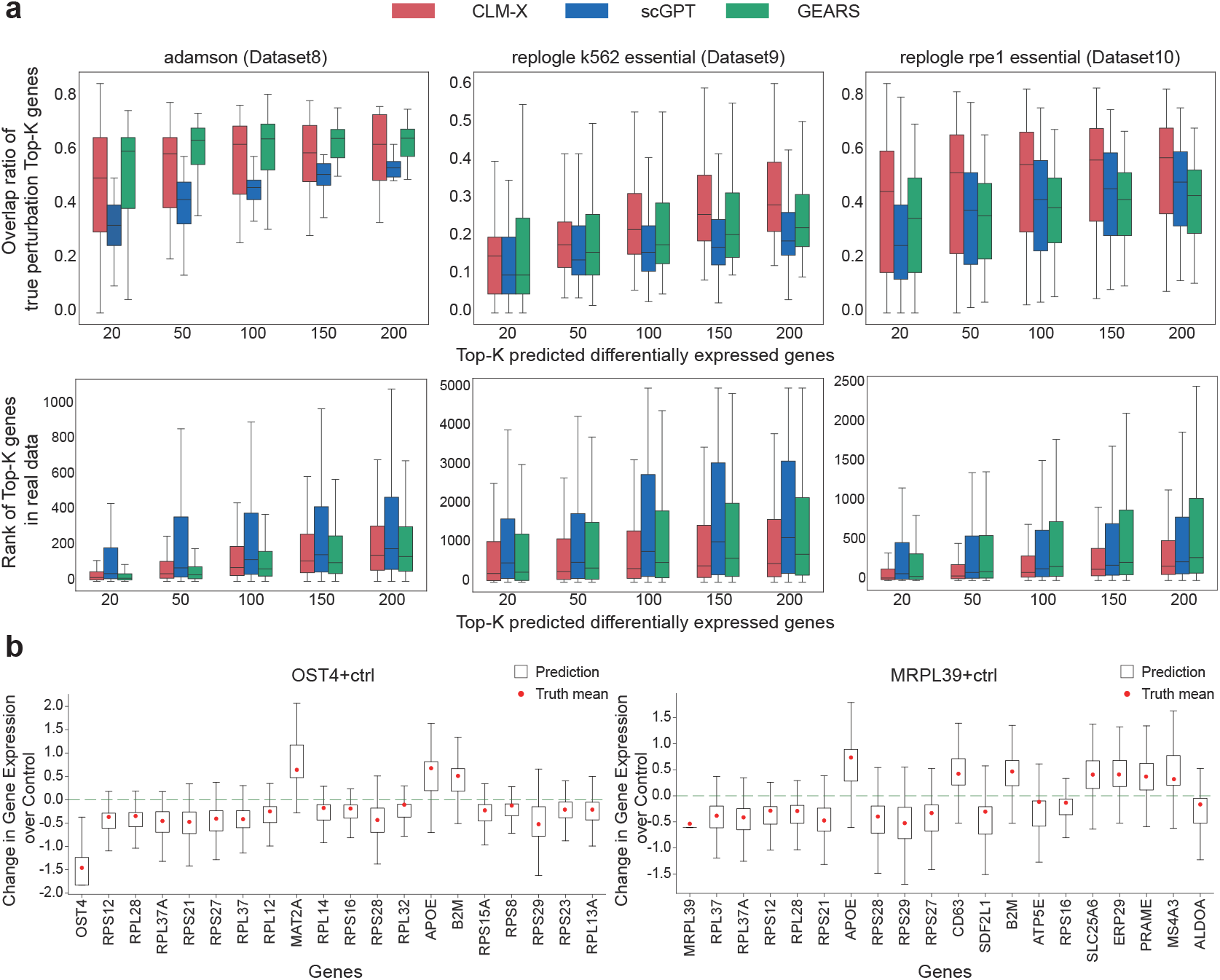
Perturbation-response prediction with CLM-X. **a**, Benchmark results on three Perturb-seq datasets (Adamson; Replogle K562 Essential; Replogle RPE1 Essential) for predicting gene-expression responses to unseen single-gene perturbations. For each dataset and method (CLM-X, scGPT, and GEARS), boxplots summarize (top) the overlap ratio between predicted and ground-truth top-*k* DE genes and (bottom) the ranks of the ground-truth top-*k* genes within the predicted gene ranking, evaluated across multiple *k* values. **b**, Adamson case studies: predicted expression changes (relative to control) for the top 20 DE genes under two representative perturbations, shown as distributions across cells alongside the ground-truth mean change for each gene.

Overall, CLM-X generalizes better to unseen perturbations, recovering both dominant DE-gene responses and genome-wide perturbation signatures. These results suggest that the learned representation transfers effectively to perturbation-response regression and supports extrapolation to previously unseen target genes, enabling more reliable in silico screening and mechanistic interpretation.

## 3 Discussion

In this study, we introduce CLM-X, a multimodal foundation model for unified analysis of scRNA-seq and scATAC-seq data. By integrating a harmonized token–value representation, a BEiT-V3 multiway Transformer backbone, and a stage-wise masked modeling reconstruction pretraining, CLM-X addresses key challenges in multimodal single-cell analysis, including modality heterogeneity, paired-data scarcity, and the need for a single pretrained encoder that supports diverse downstream analyses. CLM-X learns multimodal inter-relation primarily through bidirectional conditional completion, rather than contrastive instance discrimination. This objective encourages both alignment and complementary information transfer across modalities, while improving robustness to ATAC sparsity and binarization. Across five downstream tasks, CLM-X consistently outperforms task-specific multimodal methods and unimodal foundation models. The most pronounced gains are observed in RNA–ATAC cross-modal translation, where CLM-X supports full-scale prediction under an unfiltered-input setting and remains stable on highly variable genes. The pretrained representation also transfers well to batch correction, multimodal integration, and cell type annotation, indicating that CLM-X captures conserved biological structure while remaining resilient to technical variation; it further generalizes to perturbation response prediction.

Several limitations remain. Fixed-length packing (2,000 tokens per modality; up to 4,000 tokens for paired inputs) compresses information and may under-represent rare features. Genome-ordered ATAC patching is scalable but may miss higher-order regulatory organization and long-range interactions. In addition, imbalanced pretraining corpora and technology-specific artifacts can introduce representation bias, and large-scale pretraining with long-context fusion remains computationally expensive.

Nonetheless, CLM-X represents a step toward generalizable multimodal foundation models for cell biology, where pretrained representations can be flexibly reused across tasks, datasets, and experimental modalities. By integrating transcriptional and epigenetic information within a unified generative framework, such models have the potential to improve not only analytical robustness but also biological discovery, enabling systematic inference of regulatory programs, cellular states, and perturbation responses.

Looking forward, this framework naturally opens opportunities to further enrich multimodal representations and extend pretraining to broader data regimes, including more diverse biological contexts, additional molecular modalities, and dynamic cellular processes. Such extensions may further enhance both the robustness and interpretability of multimodal foundation models, supporting deeper biological discovery as single-cell atlases continue to expand in scale and complexity.

## 4 Methods

### 4.1 Data Processing and Tokenization

#### Gene Expression Binning and Gene Tokens

Let *X* ^RNA^ ∈ ℝ^*N*×*G*^ denote the raw cell-by-gene count matrix with *N* cells and *G* genes. Each element 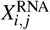 denotes the RNA abundance for gene *j ∈* {1,…, *G*} in cell *i ∈* {1,…, *N*}. Different experimental protocols and sequencing depths introduce substantial variation in absolute expression levels [40], and standard preprocessing (e.g., transcripts-per-million (TPM) normalization and the log1p transformation) does not fully remove these effects [41]. To eliminate scale while preserving within-cell ordinal information, we map nonzero counts to relative rank-preserving bins [22]. Specifically, for each cell *i* we compute *B* equal-frequency intervals 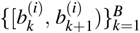 over its set of nonzero counts, with the bin edges 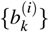estimated independently for each cell. We then define

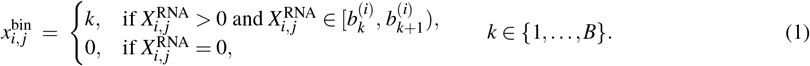

This per-cell binning makes the value semantics comparable across cells and batches (for example, 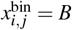 always indicates the highest within-cell expression). We set *B* = 50 during pretraining. For fine-tuning, we optionally apply the log1p transformation and highly variable gene (HVG) selection prior to binning.

We treat each gene as a token, analogous to a word in natural language. Aggregating public references and the training corpus yields a gene vocabulary of about 60k entries. The vocabulary also includes four special tokens `<cls>`, `<mask>`, `<pad>`, and `<eoc>`. For cell *i*, let *G*_*i*_ denote the number of nonzero genes. We construct a fixed-length gene-token sequence 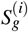 of length *L* = 2000 as follows: (1) if *G*_*i*_ ≤ *L*, list all nonzero genes in a deterministic global order and right-pad with `<pad>` to length *L*; (2) otherwise, deterministically subsample to length *L* while preserving the global order. The resulting aligned inputs are

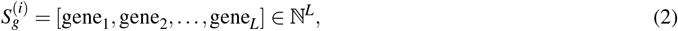

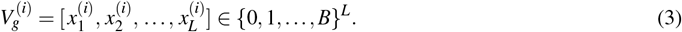

#### Chromatin Accessibility Binarization and Patch Tokens

Let *X* ^ATAC^ ∈ ℝ^*N*×*P*^ denote the cell-by-cCRE count matrix, where *N* is the number of cells and *P* is the number of candidate cis-regulatory elements (cCREs). Each entry 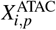 is the fragment count for cCRE *p ∈* {1,…, *P*} in cell *i ∈* {1,…, *N*}. Since most peak values are 0 or 1 and values greater than 2 are rare, we binarize all peak values to simplify their distribution and improve trainability and generalization:

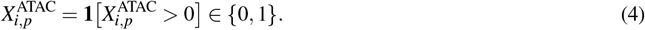

We then sort all cCREs by genomic coordinates (chromosome and start) to ensure a consistent and biologically meaningful order for downstream analysis and modeling, yielding an ordered index sequence (*p*_1_,…, *p*_*P*_). Over this ordered list, we partition 2000 contiguous, equal-capacity patches (capacity ≈575). To respect chromosome boundaries, if a patch would span two chromosomes, we drop the portion that crosses into the second chromosome. After this boundary correction, we obtain *K* = 1998 valid patch regions.

Directly treating all 1.15 million cCREs as tokens would explode the parameter count and hinder training. Therefore, we split each cell into *K* patches and treat each patch as a token to aggregate regional cCRE signals. In addition, we include four special tokens—`<cls>`, `<mask>`, `<pad>`, and `<eoc>`—to form the patch vocabulary. The token sequence for cell *i* is

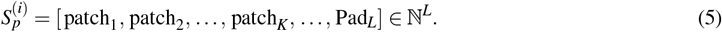

To ensure consistent inputs across patches, we standardize the length of the within-patch peak vector: if the number of peaks in a patch is smaller than a predefined unified length *M*, we pad it to length *M* (by default *M* = 600). Formally, for the *r*-th patch in cell *i*,

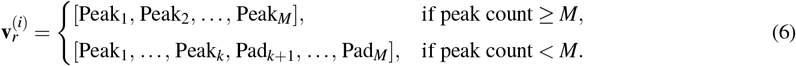

where *k* is the actual number of peaks when *k* < *M*, Peak _*j*_ *∈* {0, 1} denotes the *j*-th binarized peak within the patch, and Pad _*j*_ = 0 is the padding value. Stacking these vectors across patches yields the ATAC value vector

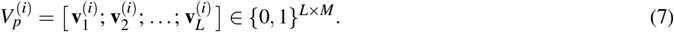

which is aligned with 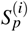 row-wise. If special tokens are prepended/appended to 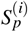, the corresponding rows in 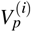 are set to the zero vector of length *M*.

### 4.2 pretraining with Multiway Transformer

#### RNA Embedding Module

The RNA embedding module contains two position-wise aligned components: a gene-identity embedding and a value embedding for expression bins. Let *d* = 512 denote the embedding dimension. The gene-identity branch is an embedding table (lookup layer) 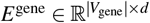. The value branch is a linear layer that maps the binned expression into *d* dimensions; we denote its output as 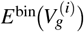. Given the aligned pair 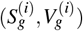 of length *L*, the RNA embedding is the position-wise sum of the two components:

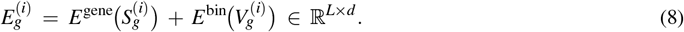

#### ATAC Embedding Module

The ATAC embedding module likewise has two aligned components: a patch-identity embedding and a peak-value embedding for the binarized accessibility vectors. The patch-identity branch is a lookup table 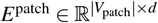. For the value branch, a linear layer projects each patch vectors 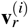 into *d* dimensions; we write the results as 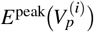. The ATAC embedding is then the position-wise sum of the two components:

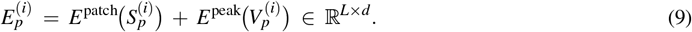

For paired RNA–ATAC inputs, we concatenate the ATAC and RNA embedding sequences along the token (sequence-length) dimension to form a single context window:

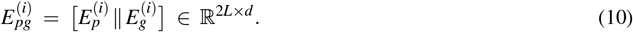

#### Multiway Transformer

CLM-X adopts a Multiway Transformer backbone in which each block couples a single shared multi-head self-attention (MHSA) module with a set of modality-specific feed-forward (FFN) experts [42, 43]. The initial inputs are

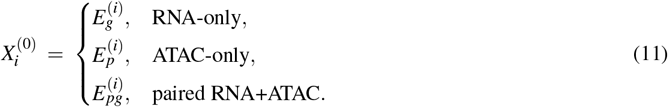

We stack *L*_blk_ pre-norm Transformer blocks (default: *L*_blk_ = 12, *H* = 8 heads). For layer *l* = 1,…, *L*_blk_:

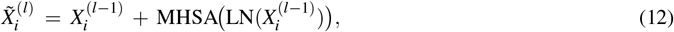

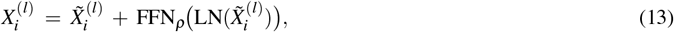

where FFN_*ρ*_ routes each token by its modality indicator ρ *∈* {R, A, RA} to the RNA expert (R-FFN), the ATAC expert (A-FFN), or the fusion expert (RA-FFN). Each expert is a two-layer MLP with GELU:

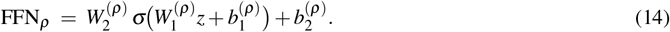

The shared MHSA is identical across input types; differences arise only from the inputs used to form (*Q, K,V*):

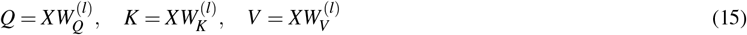

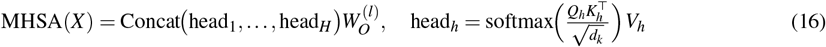

Attention computation under different inputs:

- **RNA-only**: 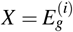; attention runs over RNA inputs only.
- **ATAC-only**: 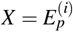; attention runs over ATAC inputs only.
- **Paired RNA+ATAC**: 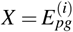; Cross-modality interactions are realized directly within the shared bidirectional attention.

#### Stage-wise pretraining

We pretrain CLM-X with a simple strategy that progressively introduces modalities while reusing (and continuously updating) the same shared MHSA parameters across all stages. The only stage-specific components are the modality-routed FFN experts and the modality-specific decoder heads. Let *L*_RNA_ and *L*_ATAC_ denote the masked reconstruction losses in Eq. (19)–(21).

1. **Stage-R (RNA-only)**. Use RNA inputs 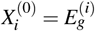, route tokens to the RNA expert (R-FFN), and optimize
2. **Stage-A (ATAC-only)**. Initialize from Stage-R, switch to ATAC inputs 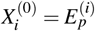, route tokens to the ATAC expert (A-FFN), and optimize *L*_ATAC_.
3. **Stage-RA (paired RNA+ATAC)**. Initialize from Stage-A, use paired inputs 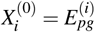, route tokens to the fusion expert (RA-FFN), and train with a two-phase schedule:

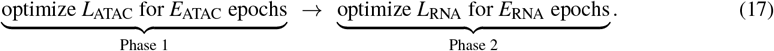

For masking, we exclude `<pad>` and special-token positions. In unimodal stages, we mask 15% of values; in the paired stage, we mask 50% of values in each modality. Given the final-layer Transformer representations, we reconstruct only the masked values. Let 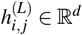 denote the final representation of gene-token position *j* in cell *i*, and 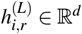 denote the final representation of patch-token position *r*. Let *M*_*R*_ be the set of masked RNA gene positions and *M*_*A*_ be the set of masked ATAC peak entries (indexed by cell *i*, patch *r*, and within-patch peak index *m*). For RNA, we regress the bin values *b*_*i, j*_ using a single-layer, fully connected neural network:

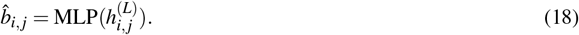

and minimize masked mean squared error

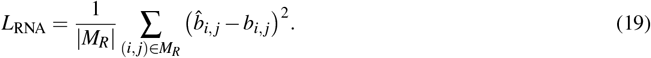

For ATAC, we use a single-layer, fully connected neural network that expands each patch embedding into *M* peak predictions. Concretely, for cell *i* and patch *r*, the decoder outputs a length-*M* logit vector

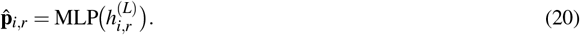

We then compute peak-wise binary cross-entropy on masked peak entries:

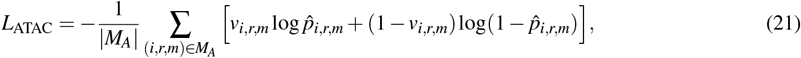

where *v*_*i,r,m*_ ∈ {*0, 1*} is the ground-truth binarized accessibility of the *m*-th peak within patch *r*.

### 4.3 Downstream Task Fine-tuning setting

#### Batch Correction

Batch effects are systematic, non-biological shifts caused by differences in experimental protocols, processing time, and sequencing depth. We fine-tune CLM-X to learn batch-aligned cell representations while preserving biological heterogeneity, following the multi-omics benchmark setting on Dataset 1–4 (PBMC, 4 batches), Dataset 5 (BMMC, 13 batches), and Dataset 6–7 (PBMC, 2 batches). For RNA, we apply library-size normalization, log1p, and Seurat HVG selection (top 2000 genes; `flavor=‘seurat’, n_top_genes=2000, min_mean=0.1, n_bins=10`). For ATAC, we binarize counts and map dataset-specific peaks to the reference cCRE space used in pretraining (*P* = 1,154,464). Let *q*_*u*_ denote a dataset-specific peak interval and *p* a reference cCRE interval; we define the overlap set

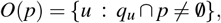

and map ATAC to the reference space by a simple “any-overlap” rule:

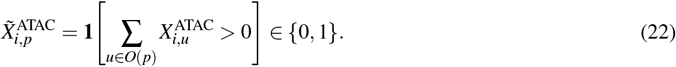

We then construct coordinate-sorted patch tokens using the same strategy as in pretraining. A `<cls>` token is prepended to aggregate cell-level information, and we use its final-layer representation 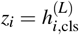 as the integrated embedding. To encourage *z*_*i*_ to capture global cell state, we attach an RNA decoder *D*_RNA_ and regress the HVG expression target 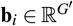 using a simple mean-squared error:

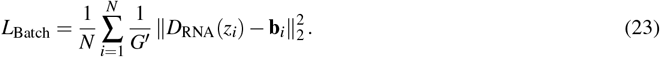

All model parameters except the newly added decoder head are initialized from the Stage-RA checkpoint, and the architecture matches pretraining (12 Transformer blocks, 8 attention heads, hidden size *d* = 512). We fine-tune on two NVIDIA A100 (80GB) GPUs with a batch size of 20 for 30 epochs. We compare against a diverse set of multimodal baselines, including deep generative integration models (MultiVI [13] and Multigrate [14]), matrix-factorization-based integration (scMoMaT [18]), and graph/topic-based integration (MIRA [15]), using benchmark-recommended protocols and hyperparameters [19]. For the zero-shot setting, we directly extract embeddings from the frozen Stage-RA checkpoint without task-specific training.

#### Multimodal Integration

Multimodal Integration aims to integrate multiple omics profiles measured from the same cells into a single representation that captures complementary biological signals across modalities. In practice, scRNA-seq reflects transcriptional programs, whereas scATAC-seq provides information about chromatin accessibility and regulatory potential; jointly modeling both modalities enables the model to learn more informative and robust cell-state features than those derived from either modality alone. We follow the same data preprocessing and input construction as in batch effect correction: RNA is library-size normalized, log1p-transformed, and restricted to Seurat-selected top-2000 HVGs, while ATAC is binarized, mapped to the reference cCRE space (*P* = 1,154,464), and converted into coordinate-sorted patch tokens. We evaluate modality fusion on Datasets 1–7. For each cell, we jointly feed the paired RNA and ATAC tokens into CLM-X, prepend a `<cls>` token, and use its final-layer representation *z*_*i*_ as the fused cell embedding; to encourage *z*_*i*_ to capture global cell state, we use the same decoder head and training objective as batch effect correction, i.e., regress the HVG expression target with the same MSE loss (Eq. 23). All model parameters except the newly added decoder head are initialized from the Stage-RA checkpoint, and the architecture matches pretraining (12 Transformer blocks, 8 attention heads, hidden size *d* = 512). We fine-tune on two NVIDIA A100 (80GB) GPUs with batch size 20 for 30 epochs, and compare against MultiVI, SCOIT, Multigrate, and MIRA under the benchmark-recommended protocols and hyperparameters[19].

#### Cross-modal Translation

Cross-modal Translation evaluates whether the model can infer an unobserved modality from an observed one, i.e., learn an explicit cross-modality mapping between RNA-seq and ATAC-seq that reflects regulatory coordination in paired multi-omics data. We fine-tune CLM-X on paired RNA–ATAC data and evaluate on Dataset 1–7 with a fixed split (80% train, 20% test). ATAC inputs are binarized and mapped to the same reference cCRE space as in pretraining (Eq. 22), then converted to patch tokens; RNA uses library-size normalized expression, and in the HVG setting, we additionally apply log1p and use Seurat top-2000 HVGs as the prediction target. We evaluate both directions: (i) ATAC →RNA under two targets: normalized full-length gene expression and log1p-transformed 2000 HVGs; (ii) RNA → ATAC, predicting reference-space accessibility from normalized full-length gene expression. For ATAC → RNA, we use the same `<cls>`-based RNA decoder as above: 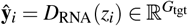, and optimize a simple averaged MSE

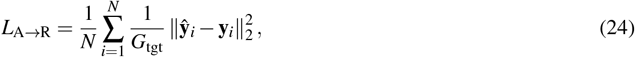

where *G*_tgt_ is either the full-gene dimension or 2000 (HVGs). For RNA ∈ ATAC, the decoder predicts a probability *â*_*i,p*_ ∈ (0, 1) for each reference cCRE *p* (equivalently obtained by flattening the patch-wise outputs back to the reference peak order). We minimize the average binary cross-entropy

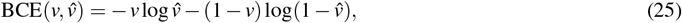

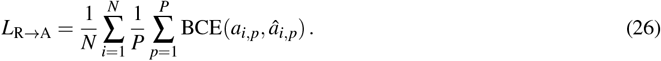

Except for the newly added decoder head(s), all parameters are initialized from Stage-RA, and the backbone configuration matches pretraining (12 blocks, 8 heads, *d* = 512). We fine-tune on two NVIDIA A100 (80GB) GPUs with batch size 20 for 100 epochs, and compare against BABEL and MultiVI following the benchmark-recommended protocol and hyperparameter settings[19].

#### Cell Type Annotation

Compared with scRNA-seq, cell type annotation for scATAC-seq is typically more challenging because accessibility profiles are sparse and high-dimensional, peak features are less directly interpretable than gene expression, and cell identity is often inferred indirectly (e.g., via motif enrichment or gene-activity surrogates). In paired RNA–ATAC settings, we formulate annotation as a fusion-based prediction problem: given both modalities for each cell, we learn a fused representation and predict cell-type labels with a supervised classifier.

For CLM-X, we evaluate three settings: CLM-X (fusion), which uses a learnable gate to adaptively reweight RNA vs. ATAC features on a per-cell basis; and two unimodal ablations, CLM-X (RNA) and CLM-X (ATAC), which use RNA-only or ATAC-only inputs, respectively. Let 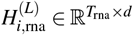 and 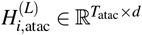 denote the final-layer hidden states of RNA and ATAC tokens for cell *i* (excluding special tokens; when both modalities are provided, these states are computed after cross-modal attention). We obtain modality-specific cell embeddings via average pooling:

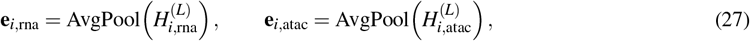

where **e**_*i*,rna_, **e**_*i*,atac_ ∈ ℝ^*d*^.

In the fusion setting, a gating network *g*(·) predicts a cell-specific fusion weight *α*_*i*_ *∈* (0, 1) from the two modality embeddings:

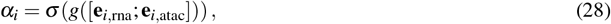

where σ (·) is the sigmoid function and [·; ·] denotes concatenation. The final fused cell representation is then computed by a differentiable weighted sum:

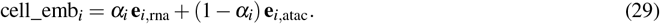

For unimodal ablations, we use the corresponding average-pooled embedding as the cell representation, i.e., cell_emb_*i*_ = **e**_*i*,rna_ for CLM-X (RNA) and cell_emb_*i*_ = **e**_*i*,atac_ for CLM-X (ATAC). The weight *#*_*i*_ and the classifier parameters are learned jointly via end-to-end training, enabling the model to automatically adjust the relative contribution of RNA and ATAC in a cell-adaptive manner.

We predict cell types with a dataset-specific classifier *C* followed by softmax, *p*_*i*_ = softmax(*C*(cell_emb_*i*_)). We optimize the standard cross-entropy loss

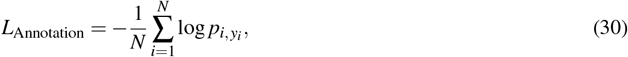

where *y*_*i*_ is the ground-truth cell type label. We fine-tune on Dataset 1–4 using an 80%/20% train/test split within each dataset. All backbone parameters are initialized from the Stage-RA checkpoint and the architecture matches pretraining (12 Transformer blocks, 8 attention heads, hidden size *d* = 512); the classifier head (and the gate network in the fusion setting) is randomly initialized to match the label set of each dataset.

Seurat (WNN) is inherently multimodal and performs annotation using a weighted-nearest-neighbor graph constructed from RNA expression and ATAC-derived features, enabling prediction from joint RNA+ATAC information. scGPT is a unimodal foundation model pretrained on scRNA-seq; to enable a fair multimodal comparison, we implement a multimodal scGPT variant that accepts paired RNA+ATAC as input in downstream fine-tuning. Specifically, we convert ATAC into a gene activity matrix and process it identically to RNA (including selecting 2,000 HVGs and applying the same tokenization procedure); the final input sequence is the concatenation of 2,000 RNA tokens and 2,000 ATAC (gene-activity) tokens. For scBridge and scJoint, we start from their official codebases and modify them to support fusion-feature annotation on paired multiome cells: both modalities are provided to produce a fused representation per cell, and a supervised classifier is trained on top of this fused representation using the same train/test splits as CLM-X. For scJoint, the original pipeline (`config.py` + `main.py`) is designed for transfer learning with labeled scRNA-seq and unlabeled scATAC-seq (gene activity) and typically outputs label-transfer results via KNN predictions (e.g., `_knn_predictions.txt`); in our fusion-annotation setting, we adapt the official implementation to use the learned representations for supervised multimodal classification rather than relying on the default KNN label-transfer output.

We fine-tune CLM-X on two NVIDIA A100 (80GB) GPUs with a batch size of 20 for 100 epochs. For baseline implementations, Seurat experiments follow Seurat v5.2.0, and scJoint experiments start from the official settings released at https://github.com/SydneyBioX/scJoint; other baseline hyperparameters follow their respective default recommendations unless otherwise stated. Since Seurat and scJoint require ATAC input as a gene activity matrix computed from raw fragments (here generated by snapATAC2), and only Dataset 1–4 in our collection can be traced back to raw fragment files, we restrict benchmarking to these four datasets.

#### Perturbation Prediction

Gene perturbation prediction aims to forecast how a genetic perturbation (e.g., CRISPR knock-out/knock-down) shifts cellular transcriptional states, which is useful for interpreting gene function, identifying regulatory dependencies, and prioritizing therapeutic targets via in silico screening. We fine-tune CLM-X to predict the post-perturbation expression profile given an unperturbed control profile together with an explicit perturbation indicator. For each dataset, we select HVGs and preprocess expression with standard normalization and log transformation for both inputs and targets; we keep the full gene vector (including zero entries) and do not apply binning/patching, using a fixed input length of *G*=5000 genes (Adamson uses *G*=5060, i.e., 5000 HVGs plus perturbation target genes). Unlike masked objectives that use different views of the same cell as input/target, we construct cross-condition pairs: each perturbed cell is randomly matched to a control cell to form an input–target pair 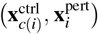. Perturbation identity is encoded by a gene-aligned categorical flag vector **s**_*i*_ ∈ {0, 1, 2} ^*G*^: for control → perturbed samples, **s**_*i*_ is 1 at the perturbed gene position and 0 elsewhere; for control → control samples (no perturbation), all entries are set to 2 as a sentinel indicating the absence of perturbation information. We add a lightweight perturbation encoder to embed **s**_*i*_ and condition the Transformer backbone, and predict the full response as

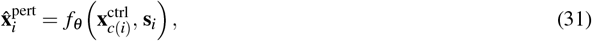

optimized with a simple averaged MSE over genes,

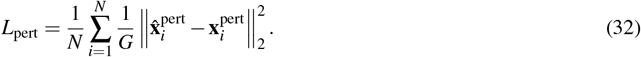

model parameters are initialized from the Stage-RA checkpoint, while the newly introduced perturbation encoder and regression head are randomly initialized. To evaluate generalization to unseen perturbations, we split train/test by perturbation condition (rather than by randomly splitting cells), ensuring that perturbation types in the test set are not observed during training. We benchmark on three single-gene perturbation datasets: Adamson (K562; 68,603 cells; 86 perturbations; 24,263 control cells; *G*=5060), replogle_k562_essential (K562; after filtering perturbations whose target genes are absent from the 5000-gene list: 68,716 cells; 412 perturbations; 10,691 control cells; *G*=5000), and replogle_rpe1_essential (RPE1; after filtering: 76,289 cells; 652 perturbations; 11,485 control cells; *G*=5000), and compare against GEARS and scGPT using their recommended perturbation-prediction settings. We fine-tune on two NVIDIA A100 (80GB) GPUs with a batch size of 20 for 100 epochs.

### 4.4 pretraining data source

#### scRNA-seq

For transcriptomic pretraining, we constructed a large-scale human scRNA-seq corpus (Human-scRNA) from the Chan Zuckerberg CELLxGENE Discover ecosystem (https://cellxgene.cziscience.com/), using CELLxGENE Census to programmatically retrieve standardized data slices in R/Python [32]. We used the Census release with schema version 2.1.0 and dataset schema version 5.2.0.

In this Census build, the Homo sapiens data comprise 1,573 datasets, with 65,601,657 cells. The associated cell-level metadata provide broad coverage of experimental and biological conditions (e.g., 31 assay categories, 827 cell types, 379 tissues / 68 tissue-general classes, 179 developmental stages, and 140 disease annotations). Leveraging this standardized metadata, we removed duplicate cells, restricted the corpus to human scRNA-seq cells, and finally obtained a Human-scRNA pretraining dataset containing approximately 36 million cells.

#### scATAC-seq

To pretrain our foundation model on single-cell chromatin accessibility profiles, we curated a dedicated scATAC-seq corpus termed Human-scATAC, integrating approximately 2.8 million scATAC-seq profiles. The dataset was built primarily from two large-scale resources: CATlas and the Descartes fetal chromatin atlas [44, 45].

CATlas (published in 2021) serves as a key pretraining source for scATAC analysis. It analyzed more than 1.3 million nuclei across 222 fetal and adult human cell types and identified roughly 1.2 million candidate cis-regulatory elements (cCREs). By integrating sci-ATAC-seq data from 30 adult tissues and 15 fetal tissues, CATlas enables fine-grained characterization of cell type-specific regulatory programs and supports systematic interpretation of non-coding variants associated with complex traits. We leveraged its scale and diversity to capture tissue- and developmental-stage-dependent chromatin accessibility dynamics during pretraining.

The Descartes fetal atlas is another indispensable component. Using sci-ATAC-seq3, it profiled chromatin accessibility in 790,957 single cells across 15 fetal organs spanning gestational weeks 8–20. The study reported 1.05 million regulatory elements, 54 TF-motif-enriched cell types, and cell type-specific heritability signals for 34 traits.

Direct integration of CATlas and Descartes required harmonization due to differences in data formats and reference genomes. Specifically, Descartes provides fragment-level data, whereas CATlas provides a cell-by-peak matrix; moreover, CATlas was processed on hg38 while Descartes fragments are in hg19 coordinates. We therefore (i) used a liftover procedure to convert Descartes coordinates from hg19 to hg38 [46], and (ii) used snapATAC2 to transform Descartes fragments into a cell-by-peak matrix consistent with CATlas [47]. During preprocessing, we additionally removed low-information cCREs to improve data quality and robustness. After these steps, the final Human-scATAC feature space contained 1,154,464 cCREs.

#### multi-omics

To introduce paired supervision during multimodal pretraining, we incorporated a previously curated pseudo-paired fetal atlas dataset, scCLIP [29]. scCLIP integrates fetal-organ scRNA-seq and scATAC-seq atlases and constructs pseudo-pairs by randomly matching cells across modalities within the same cell type, using the shared cell type annotations provided by the original studies. We used these pseudo-paired RNA–ATAC examples as paired inputs for pretraining to learn cross-modality correspondence at scale. The resulting dataset contains 377,134 pseudo-paired cells, covering 36,601 genes and 1,154,464 chromatin accessibility peaks.

### 4.5 Downstream task data source

Datasets 1–7 are paired human multiome datasets (scRNA-seq + scATAC-seq) and were used for multimodal integration and cross-modal translation. Among them, only Datasets 1–4 provide the original scATAC-seq fragments files; therefore, Datasets 1–4 were additionally used for multimodal cell type annotation to enable fair comparison with fragment-dependent baselines (Seurat (WNN), scGPT, scBridge, and scJoint). For batch correction, we formed three benchmarks: PBMC-4b (Datasets 1–4; 4 batches), BMMC-13b (Dataset 5; 13 batches), and PBMC-2b (Datasets 6–7; 2 batches). Datasets 8–10 are Perturb-seq/CRISPRi transcriptomic datasets and were used for perturbation prediction.

- **Dataset 1**. Dataset 1 is a 10x Genomics PBMC multiome (ATAC + gene expression) dataset with granulocytes removed by cell sorting (10k target recovery; 10x dataset page). We used the curated version from Hu *et al*. [19], which contains 10137 cells with paired RNA–ATAC profiles and cell type annotations. Datasets 1–4 are four public 10x PBMC multiome releases spanning two preprocessing settings (granulocyte depletion vs. no sorting) and two target sizes (10k vs. 3k), and together they define the PBMC-4b batch correction benchmark. The 10x releases provide ATAC fragments files, enabling fragment-level preprocessing when needed.
- **Dataset 2**. Dataset 2 is the corresponding 3k granulocyte-depleted PBMC multiome release from 10x Genomics (10x dataset page). The curated benchmark version contains 2592 paired profiles [19].
- **Dataset 3**. Dataset 3 is the 10x Genomics PBMC multiome dataset without cell sorting (10k target recovery; 10x dataset page). The curated benchmark version includes 8105 cells [19].
- **Dataset 4**. Dataset 4 is the unsorted 3k PBMC multiome release from 10x Genomics (10x dataset page). The curated benchmark version contains 2413 cells [19].
- **Dataset 5**. Dataset 5 is a human BMMC multiome (RNA+ATAC) dataset obtained from GEO under accession GSE194122 (https://www.ncbi.nlm.nih.gov/geo/query/acc.cgi?acc=GSE194122). It contains 69249 cells and comprises 13 batches, forming the BMMC-13b batch correction benchmark; it is also used for multimodal integration and cross-modal translation.
- **Dataset 6**. Dataset 6 is a PBMC multiome dataset accessed via GEO (GSE156478; https://www.ncbi.nlm.nih.gov/geo/query/acc.cgi?acc=GSE156478) and curated as in [48]. It contains 7468 cells and is used for multimodal integration and cross-modal translation. Together with Dataset 7, it constitutes the PBMC-2b (2-batch) batch correction benchmark.
- **Dataset 7**. Dataset 7 is the second PBMC batch from the same GEO accession (GSE156478; https://www.ncbi.nlm.nih.gov/geo/query/acc.cgi?acc=GSE156478), containing 5915 cells [48].
- **Dataset 8**. Datasets 8–10 are CRISPRi Perturb-seq transcriptomic datasets curated in [39]. Dataset 8 (Adamson) is a CRISPRi Perturb-seq dataset in the K562 leukemia cell line, containing 68603 cells with perturbation/guide annotations; the curated version used here includes 86 unique single-gene targets with matched controls.
- **Dataset 9**. Dataset 9 (Replogle_k562) is a large-scale CRISPRi Perturb-seq scRNA-seq dataset in K562 with genome-wide perturbation coverage, containing 68716 cells; each cell is associated with a perturbation label [39].
- **Dataset 10**. Dataset 10 (Replogle_rpe1) follows a similar large-scale CRISPRi Perturb-seq design profiled in the RPE1 (hTERT RPE-1) cell line, containing 162733 cells with perturbation labels [39].

### 4.6 Settings for benchmark algorithms

In this study, we benchmarked the performance of 12 algorithms across five downstream tasks (batch correction, multimodal integration, cross-modal translation, cell type annotation, and perturbation prediction). For scGPT, we extended the official codebase with multimodal support to enable ATAC inputs. For scJoint and scBridge, we extended the official implementations to perform cell type annotation using multimodal features.

- **BABEL** [36]: We used the official BABEL implementation (https://github.com/wukevin/babel) and the DANCE wrapper script (https://github.com/OmicsML/dance/blob/main/examples/multi_modality/predict_modality/babel.py), following the setup in [19]. Predictions were generated via the `predict` function of `BabelWrapper`. BABEL was used in our cross-modal translation experiments.
- **MultiVI** [13]: We used the implementation in scvi-tools (https://github.com/scverse/scvi-tools) and followed the official tutorial notebooks (https://github.com/scverse/scvi-tutorials), consistent with [19]. For modality imputation / translation, we used get_normalized_expression() and get_normalized_accessibility(). MultiVI was evaluated on batch correction, multimodal integration, and cross-modal translation.
- **Seurat (WNN)**: We used Seurat (https://github.com/satijalab/seurat). For label transfer, we ran `FindTransferAnchors(reduction = “cca”)` and `TransferData`; for multimodal feature fusion, we followed the WNN workflow (e.g., `FindMultiModalNeighbors`). Seurat (WNN) was used for the cell type annotation task.
- **Multigrate** [14]: We used the official Multigrate codebase (https://github.com/theislab/multigrate) and followed the settings in [19]. When modeling RNA with raw counts, we used `nb` loss; for log-normalized ATAC (and CLR-normalized ADT when applicable), we used `mse` loss, consistent with the recommended settings. Multigrate was evaluated on batch correction and multimodal integration.
- **scMoMaT** [18]: We used the official implementation (https://github.com/PeterZZQ/scMoMaT) and followed the preprocessing and default hyperparameters used in [19] to obtain integrated cell embeddings. scMoMaT was evaluated on batch correction.
- **MIRA** [15]: We used the official implementation (https://github.com/cistrome/MIRA) and followed the benchmark settings in [19]. For topic-model hyperparameter selection, we used `mira.topics.gradient_tune` and `model.get_learning_rate_bounds` as recommended. MIRA was evaluated on batch correction and multimodal integration.
- **CMAE** [37]: We used the authors’ official implementation of cross-modal autoencoders (https://github.com/uhlerlab/cross-modal-autoencoders) and followed the training/inference pipeline in [19]. CMAE was used in our cross-modal translation experiments.
- **SCOIT** [35]: We used the official implementation (https://github.com/deepomicslab/SCOIT) and followed the settings in [19], using the provided APIs (`fit`/`fit_list`) for model training and inference. SCOIT was evaluated on multimodal integration.
- **scJoint**: We used the official implementation (https://github.com/SydneyBioX/scJoint) and the default training configuration in `config.py`. For cell type annotation, we implemented a multimodal variant on top of the official code to leverage multimodal features. scJoint was used for the cell type annotation task.
- **scBridge**: We used the official implementation (https://github.com/XLearning-SCU/scBridge). For cell type annotation, we implemented a multimodal variant on top of the official code to leverage multimodal features. scBridge was used for the cell type annotation task.
- **scGPT** [22]: We used the official scGPT codebase and pretrained checkpoints (https://github.com/bowang-lab/scGPT). For cell type annotation, we extended the official implementation with multimodal support to enable ATAC inputs and multimodal feature fusion. For perturbation prediction, we followed the official training/evaluation scripts and fine-tuned task-specific heads when applicable. scGPT was evaluated on cell type annotation and perturbation prediction.
- **GEARS** [39]: For perturbation-response prediction, we used the official GEARS implementation (https://github.com/snap-stanford/GEARS) with its default training/evaluation pipeline. GEARS was evaluated on perturbation prediction.

### 4.7 Benchmark metrics

1. **Pearson Correlation Coefficient (PCC):** The PCC is defined as

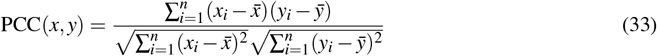

where *x*_*i*_ and *y*_*i*_ represent the abundance of protein (or chromatin accessibility at peak *i*) in cell *x* and cell *y*, respectively. 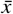 and 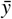 are the mean values of {*x*_*i*_} and {*y*_*i*_}, respectively. For protein–protein or peak–peak PCC, *x*_*i*_ and *y*_*i*_ denote the abundance in cell *i* for two proteins or peaks *x* and *y*.
2. **Root Mean Squared Error (RMSE) and Adjusted Root Mean Squared Error (Adj. RMSE):** We first compute the raw RMSE between predicted values **X** and true values 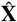:

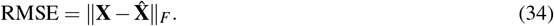

Both predicted and true values are normalized and z-scored before calculation. Since a smaller RMSE indicates better performance, for radar-chart visualization we transform RMSE into a higher-is-better score by applying min–max scaling (computed globally across all models and datasets for the corresponding task/direction) followed by inversion:

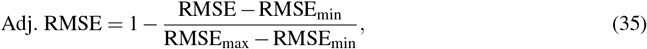

where RMSE _min_ and RMSE _max_ denote the minimum and maximum RMSE values used for scaling. After this transformation, Adj. RMSE ∈ [0, 1] and larger values indicate better performance (lower raw RMSE).
3. **Area Under Receiver Operating Characteristic Curve (AUROC)[49]:** AUROC is used to evaluate an algorithm’s ability to distinguish between binary categories (1 and 0). AUROC values range from 0 (worst) to 1 (perfect prediction); 0.5 indicates random guessing.
4. **Adjusted Rand Index (ARI)[50]e and Normalized Mutual Information (NMI)[51]:** To assess the concordance between known cell type labels and clusters identified by the Leiden algorithm, ARI and NMI are computed. Leiden clustering is performed at resolutions from 0.1 to 2.0 in increments of 0.1, and the highest ARI/NMI values are used for performance comparison.
5. **Average Silhouette Width (ASW)[52]:** ASW measures the consistency of cell–cell distances. The silhouette width for each cell is computed as

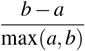

where *a* is the average intra-cluster distance and *b* is the average nearest-cluster distance. The overall ASW ranges from − 1 to 1. We use cell type ASW (cASW) and batch ASW (bASW), with linear transformations to ensure higher values indicate better performance for both metrics.
6. **Local Inverse Simpson’s Index (LISI)[53]:** LISI evaluates cell type separation (*c*LISI) and batch mixing (*i*LISI). Lower *c*LISI indicates better cell type separation; higher *i*LISI indicates better batch mixing. Linear transformations are applied so that higher values consistently indicate better performance.
7. **KNN Graph Connectivity[54]:** Connectivity is defined as

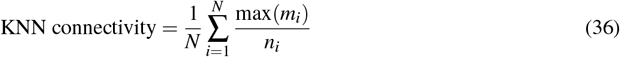

where *N* is the number of cell types, *n*_*i*_ is the number of cells in type *i*, and max(*m*_*i*_) is the size of the largest connected subgroup of type *i* in the KNN graph.
8. **Principal Component Regression (PCR)[54]:** PCR measures batch effect removal (BER) as

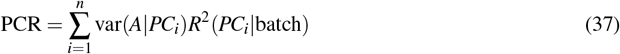

where *A* is the data matrix (RNA, ATAC, protein, or embedding), *PC*_*i*_ is the *i*-th principal component, var(*A*|*PC*_*i*_) is the variance of *A* on *PC*_*i*_, and *R*^2^(*PC*_*i*_|batch) is the squared correlation with batch labels.
9. **kBET[54]:** kBET quantifies batch effect removal by comparing batch label composition in each cell’s *k* nearest neighbors (KNN) with the overall batch composition (batch). An ideal kBET value is 1, indicating perfect removal of batch effects. We used scIB with default settings for kBET calculation.
10. **Isolated Label Score (ILS):** The ILS metric was used to evaluate the effectiveness of horizontal or mosaic integration algorithms in embedding cell connectivity graphs into low-dimensional space and isolating rare cell types present only in a subset of batches. For each cell type *i* that occurs in *k*_*i*_ batches, the ILS is defined as the average ASW value for cell types present in *k*_min_ batches, where *k*_min_ is the smallest value among all *k*_*i*_.

