## Supplementary Table for "CLM-X: A multimodal single-cell foundation model with flexible multi-way Transformer for unified scRNA-seq and scATAC-seq analysis"

### A Supplementary Figures

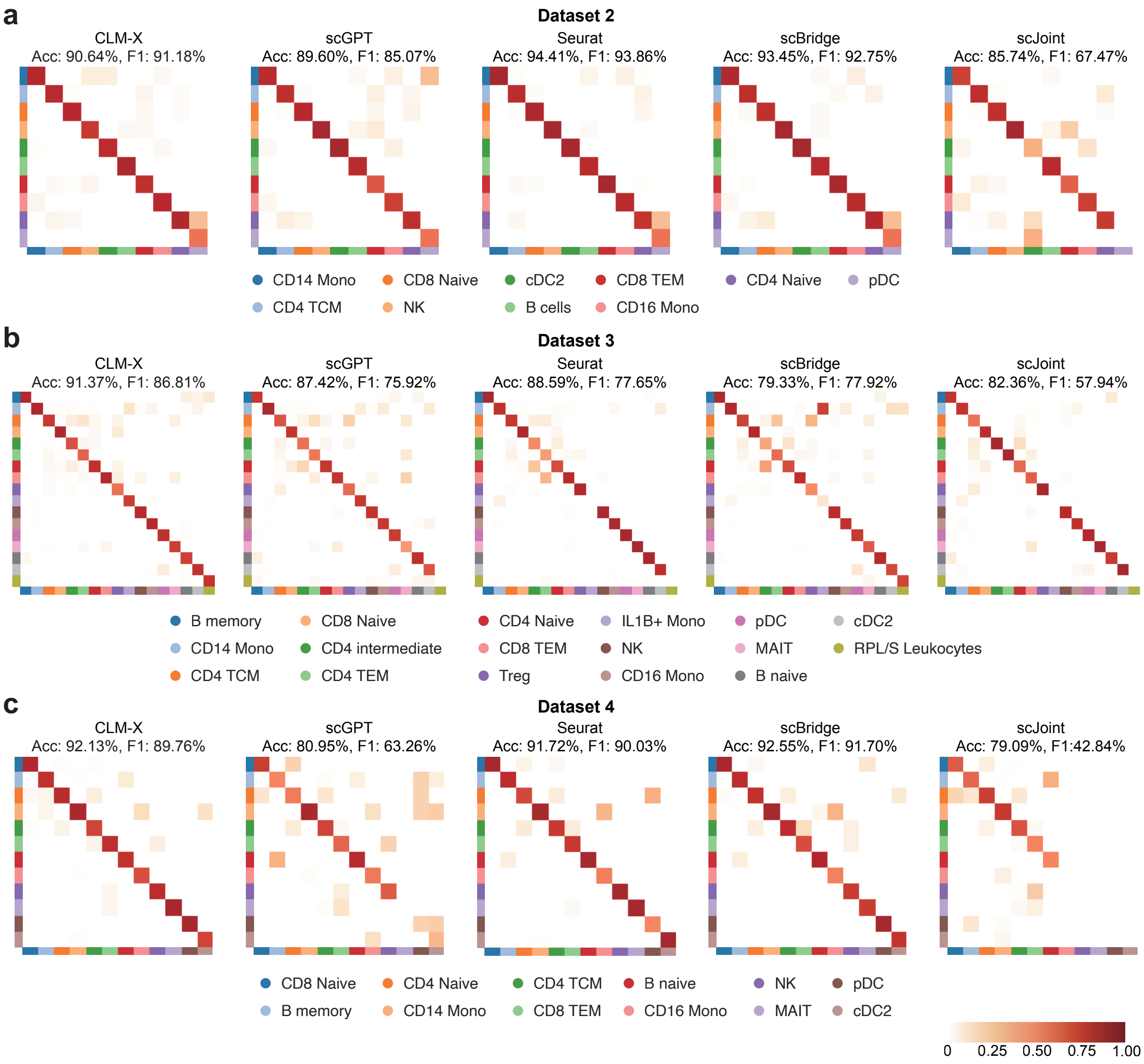


Supplementary Figure S1: Cell Type annotation result on dataset 2-4.

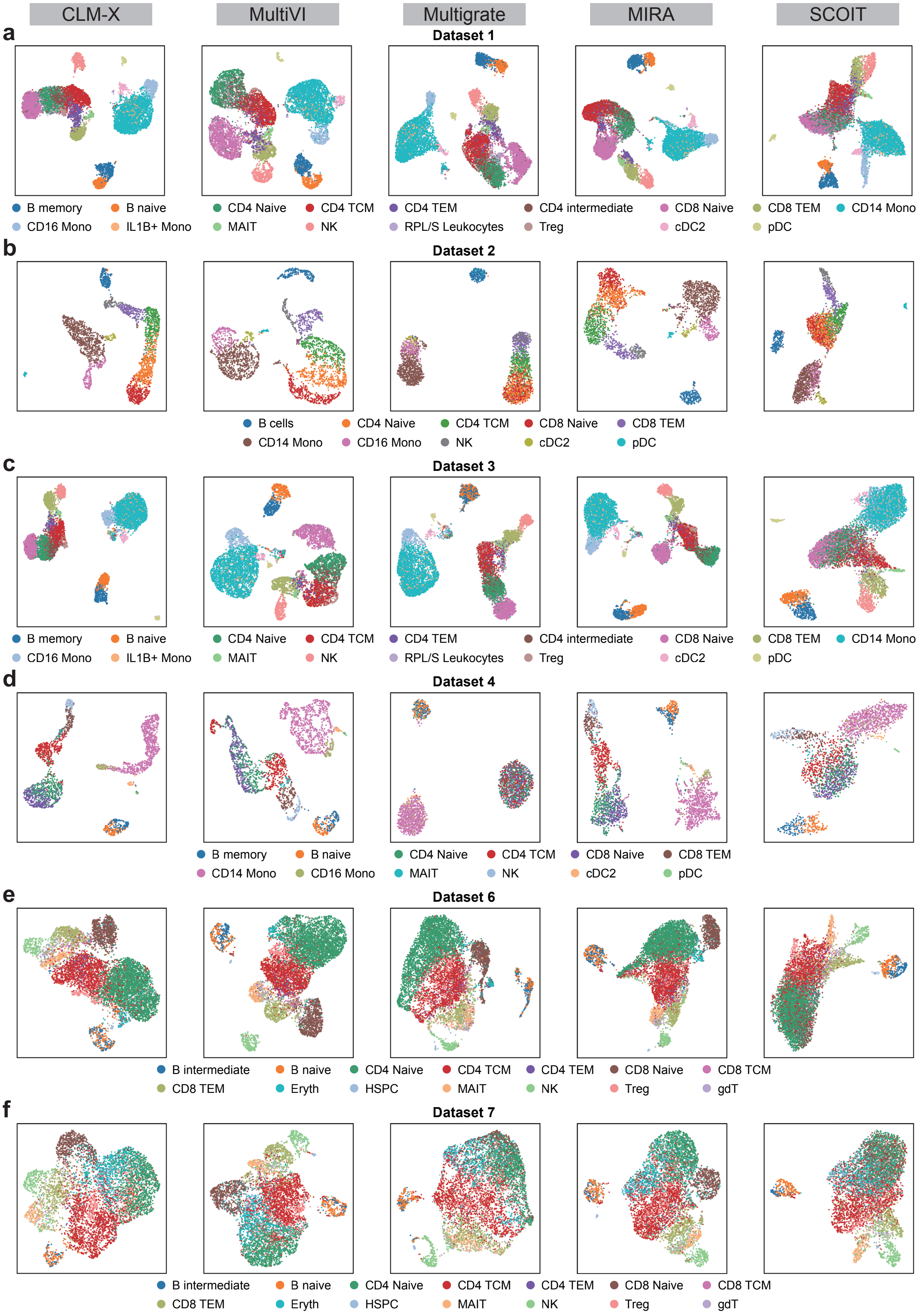


Supplementary Figure S2: Fusion UMAP on dataset 1-4, 6-7.

### B Supplementary Tables

Supplementary Table S1: **Batch-effect correction on three multiome integration benchmarks.** Results on PBMC multiome (Datasets 1–4; 4 batches; including granulocyte-depleted and unsorted samples), PBMC multiome (Datasets 6–7; 2 batches), and BMMC multiome (Dataset 5; 13 batches; multi-donor and cross-center). NMI quantifies preservation of biological structure and bASW quantifies batch mixing; Overall = $(\text{NMI} + \text{bASW})/2$. The bottom block reports mean ± SD across the three benchmarks.

| Method | NMI (↑) | bASW (↑) | Overall (↑) |
|---|---|---|---|
| **PBMC multiome (Datasets 1–4; 4 batches)** | | | |
| **CLM-X (fine-tuned)** | **0.759** | **0.972** | **0.865** |
| Multigrate | 0.734 | 0.927 | 0.831 |
| scGPT | 0.697 | 0.902 | 0.799 |
| CLM-X (zero-shot) | 0.505 | 0.884 | 0.694 |
| scMoMaT | 0.437 | 0.935 | 0.686 |
| MIRA | 0.626 | 0.482 | 0.554 |
| MultiVI | 0.547 | 0.557 | 0.552 |
| **PBMC multiome (Datasets 6–7; 2 batches)** | | | |
| **CLM-X (fine-tuned)** | **0.566** | **0.949** | **0.757** |
| Multigrate | 0.523 | 0.894 | 0.709 |
| MIRA | 0.558 | 0.851 | 0.704 |
| MultiVI | 0.514 | 0.839 | 0.677 |
| scGPT | 0.457 | 0.872 | 0.664 |
| scMoMaT | 0.342 | 0.905 | 0.623 |
| CLM-X (zero-shot) | 0.313 | 0.912 | 0.613 |
| **BMMC multiome (Dataset 5; 13 batches)** | | | |
| **CLM-X (fine-tuned)** | **0.768** | **0.973** | **0.871** |
| MIRA | 0.752 | 0.898 | 0.825 |
| Multigrate | 0.719 | 0.911 | 0.815 |
| MultiVI | 0.733 | 0.866 | 0.799 |
| scGPT | 0.676 | 0.870 | 0.773 |
| CLM-X (zero-shot) | 0.521 | 0.880 | 0.701 |
| scMoMaT | 0.189 | 0.886 | 0.538 |
| **Mean ± SD across three benchmarks** | | | |
| **CLM-X (fine-tuned)** | **0.698 ± 0.114** | **0.965 ± 0.014** | **0.831 ± 0.064** |
| Multigrate | 0.659 ± 0.118 | 0.911 ± 0.017 | 0.785 ± 0.066 |
| scGPT | 0.610 ± 0.133 | 0.881 ± 0.018 | 0.745 ± 0.072 |
| MIRA | 0.645 ± 0.098 | 0.744 ± 0.228 | 0.694 ± 0.136 |
| MultiVI | 0.598 ± 0.118 | 0.754 ± 0.171 | 0.676 ± 0.124 |
| CLM-X (zero-shot) | 0.446 ± 0.116 | 0.892 ± 0.017 | 0.669 ± 0.049 |
| scMoMaT | 0.323 ± 0.125 | 0.909 ± 0.025 | 0.616 ± 0.074 |

Supplementary Table S2: **Integration performance on seven multiome benchmarks (Datasets 1–7).** Fusion integrates paired scRNA-seq and scATAC-seq profiles into a unified cell embedding across six PBMC multiome datasets (Datasets 1–4, 6–7) and one BMMC multiome dataset (Dataset 5). ARI and NMI quantify agreement between clustering assignments and cell-type labels; cLISI quantifies local consistency of cell-type neighborhoods; cASW quantifies cell-type separability. Overall is computed as $(\mathrm{ARI}+\mathrm{NMI}+\mathrm{cLISI}+\mathrm{cASW})/4$. The bottom block reports mean ± SD across datasets.

| Method | ARI (↑) | NMI (↑) | cLISI (↑) | cASW (↑) | Overall (↑) |
|---|---|---|---|---|---|
| **Dataset 1** | | | | | |
| CLM-X | **0.769** | **0.795** | 0.988 | 0.551 | 0.776 |
| MultiVI | 0.765 | 0.786 | **0.992** | **0.586** | **0.782** |
| Multigrate | 0.743 | 0.772 | 0.987 | 0.545 | 0.762 |
| MIRA | 0.745 | 0.763 | 0.988 | 0.558 | 0.764 |
| SCOIT | 0.551 | 0.677 | 0.969 | 0.523 | 0.680 |
| **Dataset 2** | | | | | |
| CLM-X | 0.703 | 0.800 | **0.992** | **0.701** | 0.799 |
| MultiVI | **0.785** | **0.813** | 0.987 | 0.635 | **0.805** |
| Multigrate | 0.590 | 0.702 | 0.904 | 0.524 | 0.680 |
| MIRA | 0.643 | 0.734 | 0.978 | 0.614 | 0.742 |
| SCOIT | 0.544 | 0.693 | 0.918 | 0.512 | 0.667 |
| **Dataset 3** | | | | | |
| CLM-X | **0.762** | **0.772** | 0.984 | 0.541 | **0.765** |
| MultiVI | 0.601 | 0.732 | **0.991** | **0.586** | 0.727 |
| Multigrate | 0.601 | 0.733 | 0.978 | 0.528 | 0.710 |
| MIRA | 0.741 | 0.746 | 0.989 | 0.554 | 0.757 |
| SCOIT | 0.571 | 0.644 | 0.958 | 0.506 | 0.670 |
| **Dataset 4** | | | | | |
| CLM-X | 0.585 | **0.749** | **0.974** | **0.618** | **0.731** |
| MultiVI | 0.630 | 0.676 | 0.973 | 0.569 | 0.712 |
| Multigrate | 0.439 | 0.614 | 0.846 | 0.495 | 0.598 |
| MIRA | **0.643** | 0.715 | 0.954 | 0.550 | 0.715 |
| SCOIT | 0.546 | 0.655 | 0.923 | 0.486 | 0.652 |
| **Dataset 5** | | | | | |
| CLM-X | **0.718** | **0.785** | 0.988 | 0.544 | **0.759** |
| MultiVI | 0.540 | 0.729 | **0.991** | **0.586** | 0.711 |
| Multigrate | 0.696 | 0.782 | 0.983 | 0.574 | **0.759** |
| MIRA | 0.000 | 0.014 | 0.729 | 0.470 | 0.303 |
| SCOIT | 0.507 | 0.650 | 0.955 | 0.508 | 0.655 |
| **Dataset 6** | | | | | |
| CLM-X | 0.608 | 0.583 | 0.968 | 0.523 | 0.671 |
| MultiVI | **0.614** | **0.618** | **0.977** | **0.559** | **0.692** |
| Multigrate | 0.481 | 0.560 | 0.967 | 0.528 | 0.634 |
| MIRA | 0.594 | 0.566 | 0.969 | 0.516 | 0.661 |
| SCOIT | 0.389 | 0.474 | 0.929 | 0.496 | 0.572 |
| **Dataset 7** | | | | | |
| CLM-X | **0.464** | **0.529** | **0.942** | 0.505 | **0.610** |
| MultiVI | 0.419 | 0.497 | 0.935 | **0.517** | 0.592 |
| Multigrate | 0.269 | 0.405 | 0.901 | 0.482 | 0.514 |
| MIRA | 0.425 | 0.495 | 0.927 | 0.510 | 0.589 |
| SCOIT | 0.241 | 0.397 | 0.881 | 0.450 | 0.492 |
| **Mean ± SD across Datasets 1–7** | | | | | |
| **CLM-X** | **0.658 ± 0.111** | **0.716 ± 0.112** | 0.977 ± 0.017 | 0.569 ± 0.068 | **0.730 ± 0.067** |
| MultiVI | 0.622 ± 0.126 | 0.693 ± 0.108 | **0.978 ± 0.020** | **0.577 ± 0.036** | 0.717 ± 0.069 |
| Multigrate | 0.546 ± 0.163 | 0.653 ± 0.136 | 0.938 ± 0.055 | 0.525 ± 0.030 | 0.665 ± 0.090 |
| MIRA | 0.542 ± 0.262 | 0.576 ± 0.268 | 0.933 ± 0.093 | 0.539 ± 0.046 | 0.647 ± 0.164 |
| SCOIT | 0.478 ± 0.121 | 0.599 ± 0.115 | 0.933 ± 0.030 | 0.497 ± 0.024 | 0.627 ± 0.069 |

Supplementary Table S3: **Cross-modality translation performance on seven multiome benchmarks (Datasets 1–7).** We evaluate RNA→ATAC translation over all peaks using AUROC (↑), Pearson correlation coefficient (PCC; ↑), and root mean square error (RMSE; ↓). We also evaluate ATAC→RNA translation using either normalized all-gene targets or log-transformed 2,000 highly variable genes (HVGs), reporting PCC (↑) and RMSE (↓). Best values are highlighted in bold for each dataset; the bottom block reports mean ± SD across Datasets 1–7.

| | RNA→ATAC (all peaks) | | | ATAC→RNA (norm all genes) | | ATAC→RNA (log 2,000 HVGs) | |
|---|---|---|---|---|---|---|---|
| Method | AUROC (↑) | PCC (↑) | RMSE (↓) | PCC (↑) | RMSE (↓) | PCC (↑) | RMSE (↓) |
| **Dataset 1** | | | | | | | |
| CLM-X | 0.938 | 0.444 | **0.088** | **0.920** | **1.400** | **0.686** | **0.352** |
| BABEL | 0.651 | 0.309 | 0.094 | 0.910 | 1.532 | 0.601 | 0.382 |
| MultiVI | **0.949** | **0.449** | 0.102 | 0.885 | 3.576 | 0.603 | 0.500 |
| CMAE | 0.633 | 0.262 | 0.096 | 0.893 | 1.839 | 0.495 | 0.432 |
| **Dataset 2** | | | | | | | |
| CLM-X | 0.928 | **0.409** | **0.083** | **0.890** | **1.604** | **0.687** | **0.343** |
| BABEL | 0.642 | 0.285 | 0.088 | 0.883 | 1.726 | 0.598 | 0.375 |
| MultiVI | **0.942** | 0.397 | 0.117 | 0.662 | 3.524 | 0.519 | 0.487 |
| CMAE | 0.568 | 0.215 | 0.092 | 0.798 | 2.588 | 0.462 | 0.434 |
| **Dataset 3** | | | | | | | |
| CLM-X | 0.984 | **0.479** | **0.067** | 0.898 | **1.670** | **0.620** | **0.361** |
| BABEL | 0.672 | 0.318 | 0.071 | **0.906** | 1.695 | 0.561 | 0.386 |
| MultiVI | **0.989** | 0.470 | 0.085 | 0.836 | 3.798 | 0.505 | 0.473 |
| CMAE | 0.581 | 0.189 | 0.074 | 0.877 | 2.101 | 0.436 | 0.436 |
| **Dataset 4** | | | | | | | |
| CLM-X | 0.921 | **0.352** | **0.073** | **0.857** | **1.982** | **0.642** | **0.352** |
| BABEL | 0.660 | 0.257 | 0.074 | 0.852 | 2.099 | 0.614 | 0.365 |
| MultiVI | **0.939** | 0.326 | 0.122 | 0.573 | 3.865 | 0.467 | 0.471 |
| CMAE | 0.531 | 0.057 | 0.076 | 0.752 | 2.844 | 0.382 | 0.438 |
| **Dataset 5** | | | | | | | |
| CLM-X | 0.981 | **0.395** | **0.054** | **0.892** | **4.816** | **0.529** | **0.493** |
| BABEL | 0.669 | 0.261 | 0.057 | 0.753 | 6.766 | 0.486 | 0.507 |
| MultiVI | **0.988** | 0.378 | 0.084 | 0.671 | 11.788 | 0.349 | 0.601 |
| CMAE | 0.560 | 0.030 | 0.060 | 0.694 | 10.942 | 0.298 | 0.565 |
| **Dataset 6** | | | | | | | |
| CLM-X | 0.983 | **0.540** | **0.095** | **0.809** | **1.242** | **0.629** | **0.402** |
| BABEL | 0.730 | 0.361 | 0.105 | 0.509 | 1.845 | 0.610 | 0.411 |
| MultiVI | **0.986** | 0.501 | 0.129 | 0.145 | 2.138 | 0.507 | 0.536 |
| CMAE | 0.764 | 0.379 | 0.108 | 0.773 | 1.455 | 0.493 | 0.480 |
| **Dataset 7** | | | | | | | |
| CLM-X | 0.980 | **0.546** | **0.101** | **0.754** | **1.346** | **0.624** | **0.392** |
| BABEL | 0.733 | 0.371 | 0.110 | 0.520 | 1.783 | 0.615 | 0.398 |
| MultiVI | **0.986** | 0.512 | 0.135 | 0.668 | 2.074 | 0.541 | 0.519 |
| CMAE | 0.598 | 0.258 | 0.117 | 0.721 | 1.559 | 0.488 | 0.457 |
| **Mean ± SD across Datasets 1–7** | | | | | | | |
| **CLM-X** | 0.959 ± 0.029 | **0.452 ± 0.074** | **0.080 ± 0.016** | **0.860 ± 0.059** | **2.009 ± 1.262** | **0.631 ± 0.053** | **0.385 ± 0.052** |
| BABEL | 0.680 ± 0.037 | 0.309 ± 0.045 | 0.086 ± 0.019 | 0.762 ± 0.177 | 2.492 ± 1.892 | 0.584 ± 0.047 | 0.403 ± 0.048 |
| MultiVI | **0.968 ± 0.024** | 0.433 ± 0.069 | 0.111 ± 0.021 | 0.634 ± 0.242 | 4.395 ± 3.347 | 0.499 ± 0.078 | 0.512 ± 0.046 |
| CMAE | 0.605 ± 0.077 | 0.199 ± 0.122 | 0.089 ± 0.020 | 0.787 ± 0.075 | 3.333 ± 3.394 | 0.436 ± 0.073 | 0.463 ± 0.048 |

Supplementary Table S4: **Multimodal cell type annotation on four matched 10x Genomics PBMC multiome datasets (Datasets 1–4).** Within each dataset, we perform an 80/20 stratified cell-level split by cell-type labels and report Accuracy and Macro F1 (both in %; ↑ indicates higher is better). CLM-X is supervised fine-tuned for cell type classification using fused (RNA+ATAC) cell embeddings; we additionally report unimodal CLM-X variants and external baselines (Seurat (WNN), scGPT, scBridge, scJoint). Best results are bolded for each dataset and metric; the bottom block reports mean ± s.d. across Datasets 1–4.

| Method | Accuracy (%, ↑) | Macro F1 (%, ↑) |
|---|---|---|
| **Dataset 1** | | |
| CLM-X (Fusion) | **90.38** | **85.22** |
| CLM-X (RNA) | 89.55 | 84.65 |
| CLM-X (ATAC) | 80.77 | 67.27 |
| scGPT | 87.48 | 75.49 |
| Seurat (WNN) | 89.10 | 79.80 |
| scBridge | 80.18 | 76.42 |
| scJoint | 83.38 | 64.31 |
| **Dataset 2** | | |
| CLM-X (Fusion) | 93.64 | 91.18 |
| CLM-X (RNA) | 88.85 | 89.06 |
| CLM-X (ATAC) | 83.65 | 81.73 |
| scGPT | 89.60 | 85.07 |
| Seurat (WNN) | **94.41** | **93.86** |
| scBridge | 93.44 | 92.75 |
| scJoint | 85.74 | 67.47 |
| **Dataset 3** | | |
| CLM-X (Fusion) | **91.37** | **86.81** |
| CLM-X (RNA) | 90.33 | 85.68 |
| CLM-X (ATAC) | 75.77 | 57.14 |
| scGPT | 87.42 | 75.92 |
| Seurat (WNN) | 89.20 | 78.08 |
| scBridge | 79.33 | 77.92 |
| scJoint | 82.36 | 57.94 |
| **Dataset 4** | | |
| CLM-X (Fusion) | 92.13 | 89.76 |
| CLM-X (RNA) | 91.32 | 88.07 |
| CLM-X (ATAC) | 77.69 | 63.65 |
| scGPT | 80.59 | 63.26 |
| Seurat (WNN) | 91.72 | 90.01 |
| scBridge | **92.55** | **91.70** |
| scJoint | 79.09 | 42.84 |
| **Mean ± s.d. across Datasets 1–4** | | |
| **CLM-X (Fusion)** | **91.88 ± 1.19** | **88.24 ± 2.35** |
| Seurat (WNN) | 91.11 ± 2.18 | 85.44 ± 6.67 |
| CLM-X (RNA) | 90.01 ± 0.92 | 86.86 ± 1.77 |
| scBridge | 86.38 ± 6.63 | 84.70 ± 7.56 |
| scGPT | 86.27 ± 3.40 | 74.94 ± 7.75 |
| scJoint | 82.64 ± 2.39 | 58.14 ± 9.48 |
| CLM-X (ATAC) | 79.47 ± 3.00 | 67.45 ± 9.01 |

Supplementary Table S5: **Dataset 1 class-wise performance on IL1B+ Mono and RPL/S Leukocytes.** Recall is computed as diagonal/row-sum (per-class accuracy), and precision as diagonal/column-sum (label purity), from the confusion matrices in Fig. 5c.

| | IL1B+ Mono | | RPL/S Leukocytes | |
|---|---|---|---|---|
| Method | Recall (%, ↑) | Precision (%, ↑) | Recall (%, ↑) | Precision (%, ↑) |
| CLM-X | 82.35 | 57.14 | 39.29 | 73.33 |
| scBridge | 70.59 | 15.09 | 39.29 | 28.21 |
| Seurat | 0.00 | 0.00 | 3.57 | 25.00 |
| scGPT | 0.00 | 0.00 | 0.00 | 0.00 |
| scJoint | 0.00 | 0.00 | 0.00 | 0.00 |

Supplementary Table S6: **Pearson correlation of predicted vs. ground-truth perturbation-induced differential expression under a held-out perturbation split on three Perturb-seq benchmarks.** We use $\Delta$ to denote the ground-truth differential expression vector (perturbation vs. control) and $\widehat{\Delta}$ its prediction. **DE20 Pearson** is the Pearson correlation between $\widehat{\Delta}$ and $\Delta$ restricted to the top-20 ground-truth DE genes; $\Delta$ **Pearson** is the Pearson correlation computed over the full gene set (higher is better, ↑). Results are reported on Adamson , Replogle K562 Essential , and Replogle RPE1 Essential.

| Method | DE20 Pearson (↑) | $\Delta$ Pearson (↑) |
|---|---|---|
| **Adamson** | | |
| GEARS | 0.9659 | **0.7467** |
| scGPT | 0.9695 | 0.6192 |
| CLM-X | **0.9724** | 0.7423 |
| **Replogle K562 Essential** | | |
| GEARS | 0.9087 | 0.2784 |
| scGPT | 0.9331 | 0.3186 |
| CLM-X | **0.9527** | **0.3518** |
| **Replogle RPE1 Essential** | | |
| GEARS | 0.7204 | 0.5021 |
| scGPT | 0.7828 | 0.5686 |
| CLM-X | **0.9121** | **0.6123** |
